# SF3B1 phosphorylation is an evolutionarily conserved step in spliceosome activation carried out by the divergent, OTS964-insensitive kinase CRK9 in trypanosomes

**DOI:** 10.1101/2025.11.20.689580

**Authors:** Kazuya Machida, Priyanka Rattan, Nuala Peterman, Bing Hao, Arthur Günzl

## Abstract

SF3B1 is a subunit of the heptameric SF3B complex which, as part of the U2 small nuclear ribonucleoprotein, facilitates branch point recognition in pre-mRNA splicing. In addition to this early-stage function, it was recently shown that activation of the spliceosome depends on the phosphorylation of threonine-proline (TP) motifs in SF3B1’s N-terminal domain (NTD) by cyclin-dependent kinase 11 (CDK11). This breakthrough result was made possible by the discovery of the CDK11-specific inhibitor OTS964. Trypanosomes are protistan parasites whose proteomes are highly divergent in sequence from those of model organisms, and thus their CDKs were generically named CDC2-related kinases (CRKs). We previously characterized the trimeric CRK9 complex of *Trypanosoma brucei* and showed that it is essential for spliced leader *trans* splicing, the predominant splicing mode in the parasite. Although CRK9 and CDK11 deviate from each other substantially, we show that CRK9 activity is required to maintain SF3B1 phosphorylation *in vivo*, CRK9 directly phosphorylates TP motifs in the SF3B1 NTD *in vitro*, and the TP motifs themselves are crucial for spliceosome activation, demonstrating evolutionary conservation of this essential splicing step. Contrary to CDK11 and human cells, CRK9 and trypanosomes were rather insensitive to OTS964, indicating potentially exploitable differences in their ATP-binding pockets.

## Introduction

Pre-mRNA splicing is carried out by the spliceosome, a large and dynamic ribonucleoprotein assembly of five uridine-rich, small nuclear RNAs (U snRNAs) and about 85 conserved spliceosomal core proteins that are conserved between budding yeast *Saccharomyces cerevisiae* and human cells (1–3). Spliceosome assembly is a stepwise process in which, very briefly, recruitment of the U1 small nuclear ribonucleoprotein (U1 snRNP) to the 5’ splice site and of the U2 snRNP to the branch point leads to the formation of the A complex, and the subsequent recruitment of the U4/U6.U5 triple snRNP to the formation of the precatalytic B complex. For activation, i.e. the formation of the B^act^ complex, the spliceosome undergoes multiple rearrangements which includes the discard of the U1 and U4 snRNPs.

SF3B1 is a core splicing factor and a key subunit of the heptameric SF3B complex which assembles into the U2 small nuclear ribonucleoprotein (U2 snRNP) and facilitates branchpoint recognition and base-pairing of U2 snRNA with the branch site in the early stages of spliceosome assembly. It is a scaffold protein that undergoes multiple interactions with SF3B subunits and other splicing factors such as U2 Auxiliary Factor 2 (U2AF2, aka U2AF65), p14 (aka PHF5A, SF3b14b) and SUGP1, and it directly contacts the pre-mRNA on both sides of the branch site (4). The importance of SF3B1 for pre-mRNA splicing and cell function is underscored by the fact that it is the most frequently mutated spliceosomal protein in different types of cancer, and a target of anti-cancer therapy (4–7). SF3B1 has an intrinsically unfolded N-terminal domain (NTD) that contains seven U2AF Ligand Motifs (ULMs) and an SF3B6 binding motif, a large HEAT repeat domain that folds into a superhelix, and a C-terminal anchor region that directly interacts with subunits SF3B3 and SF3B5 (8). In addition to its essential function in forming the pre-catalytic spliceosome, SF3B1 was found to be phosphorylated subsequent to these early steps, specifically around spliceosome activation when the U4 snRNP is discarded from the spliceosome and the B^act^ spliceosome is formed (9).

In human cells, cyclin-dependent kinase 11 (CDK11) forms an enzyme complex with the nearly identical cyclins L1 or L2 (from here on referred to as cyclin L), and early studies suggested that its activity is linked to pre-mRNA splicing through phosphorylating the SR protein SRSF7 and directly interacting with the SR-related protein RNPS1 (10,11). However, discovery of the CDK11-specific inhibitor OTS964 (12) facilitated the unearthing of a fundamentally important role of CDK11 in pre-mRNA splicing. The kinase phosphorylates SF3B1 specifically during splicing activation, and the known phospho-sites are TP motifs in the NTD (13). Moreover, when CDK11 was blocked by OTS964, the pre-catalytic B complex accumulated and the transition to the active B complex (B^act^) was impaired, indicating that SF3B1 phosphorylation by CDK11 is an essential step in spliceosome activation (13). *S. cerevisiae* does not possess a CDK11 homolog, and in the fission yeast *Schizosaccharomyces pombe* CDK11 is encoded by a non-essential gene and indirectly regulates expression of 55 genes by phosphorylating two subunits of the mediator complex (14). Consequently, it was proposed that SF3B1 phosphorylation by CDK11 as an essential step for splicing activation has specifically evolved in metazoans (15).

Trypanosomatids comprise a phylogenetic phylum of protistan parasites that include *Trypanosoma brucei*, *T. cruzi*, and *Leishmania* spp. which cause devastating human diseases in large parts of the world (16). Trypanosomatids are distantly related to model organisms such as mammals and yeast (17), and pre-mRNA processing in these organisms has drawn attention because all their mRNAs are processed from polycistronic precursors by spliced leader (SL) *trans* splicing and polyadenylation (18,19). In SL *trans* splicing, the capped 5^/^-terminal sequence of the small nuclear SL RNA is spliced onto the 5^/^ end of every mRNA. As other eukaryotes, trypanosomatids possess introns, i.e. carry out *cis* splicing, but only three genes have been found that carry a single intron, including *PAP1* which encodes poly(A) polymerase 1 (20–23). *Trans* and *cis* splicing are mediated by the spliceosome and occur by the same two consecutive transesterifications (24,25). Trypanosome splicing factors are highly divergent in sequence to those of model organisms, but their spliceosomal protein repertoire matches that of yeast and harbors several essential, seemingly trypanosome-specific proteins (18,26).

CDKs are considered highly druggable enzymes, and several CDK inhibitors have been approved as anti-cancer drugs (27). *T. brucei* possesses 11 CDKs which are extremely divergent in sequence to CDKs of other eukaryotes and, thus, were generically named <u>C</u>DC2-<u>r</u>elated <u>k</u>inases or CRKs (28). CRK9 is encoded by an essential gene [accession number Tb927.2.4510 at TriTrypDB.org (29,30)], localized in the nucleus, and its kinase activity indispensable for pre-mRNA *trans* and *cis* splicing (31–33). CRK9 is an unusual CDK because it forms a trimeric complex with cyclin 12 (CYC12; Tb927.10.9160) and a small CRK9-associated protein (CRK9AP; Tb927.3.4170), and its kinase domain as well as both cyclin domains in CYC12 are disrupted by extensive sequence insertions (34). In addition, the CDK11-specific cyclin binding helix motif PITSLRE, which is conserved from human to *S. pombe*, is present in trypanosome CRK12 but not in CRK9. On the other hand, trypanosome CYC12 possesses a highly charged C-terminus reminiscent of the C-terminal RS domain of human cyclin L (34,35). A second parallel was the recent finding that human SAP30BP functions as CRK9AP in trypanosomes. We had shown previously that CRK9AP binding to CYC12 is required for the formation of the CRK9 complex, and that *CRK9AP* silencing led to a rapid loss of CRK9 and CYC12 protein *in vivo,* resulting in a global splicing defect (34). Accordingly, SAP30BP binding to cyclin L facilitated the assembly of the CDK11 enzyme complex, and its ablation negatively affected the stability of CDK11 and caused a general splicing defect (36).

Here we show that trypanosome SF3B1 (Tb927.11.11850) has a phosphorylated form in *T. brucei* and, by using analog-sensitive kinase technology, that trypanosome CRK9 activity is required to maintain SF3B1 phosphorylation *in vivo*, both proteins interact with each other, and CRK9 phosphorylates TP motifs in the SF3B1 NTD *in vitro*. Although specific spliceosomal complexes have not been characterized in *T. brucei*, our data strongly indicate that CRK9 inhibition interferes with spliceosome activation in the parasite, identifying SF3B1 NTD phosphorylation as an evolutionary ancient step in this process. Moreover, we found that cultured trypanosomes and CRK9 activity are magnitudes less sensitive to OTS964 than human cells and CDK11, supporting the notion that, despite the functional overlap, CRK9 may be amenable to selective inhibition.

## Materials and Methods

### DNAs

Plasmids CRK9^WT^-PTP-NEO, CRK9^AS^-PTP-NEO and PRP19-PTP were described previously (32,37). The PTP tag is a composite tag, consisting of a tandem protein A (<u>P</u>rotA) domain, a tobacco etch virus (<u>T</u>EV) cleavage site and the protein C epitope (<u>P</u>rotC) (38). Plasmid PURO-HA-SF3B1 is a derivative of pN-PURO-PTP (38) in which the PTP tag sequence was replaced by the HA tag sequence, followed by an XhoI restriction site, coding positions 4-723 of the *T. brucei brucei* 427 SF3B1 gene and a KpnI restriction site. For stable transfection of trypanosomes, the plasmid was linearized with BamHI. To generate a cell line that exclusively expressed HA-tagged, analog-sensitive CRK9, we generated pCRK9^AS^-HA-PURO by replacing the tag sequence and the selectable marker gene of CRK9^AS^-PTP-NEO. For conditional silencing of *SF3B1*, we constructed pT7-SF3B1-3^/^UTR-stl by inserting the first 370 bp of the 3^/^ UTR sequence in a sense-stuffer-antisense arrangement into the HindIII and MluI restriction sites of the pT7-stl vector (39), downstream of its doxycycline (dox)-inducible T7 promoter. For conditional transgene expression of full-length, HA-tagged SF3B1, we constructed pT7-SF3B1-HA-BLA by replacing the CITFA2 coding regions in pT7-CITFA2-HA (40) with that of SF3B1 and exchanging the bleomycin resistance gene with the blasticidin S deaminase gene. Please note that in comparison to the genome database, the cloned *SF3B1* sequence contained one nucleotide mismatch, T3104C, which resulted in a V1035A amino acid change. All pT7-stl derivatives were linearized within its ribosomal spacer target sequence with EcoRV. For recombinant expression of the GST-tagged N- and C-terminal domains of SF3B1 in *Escherichia coli*, we generated pGEX6P1-SF3B1-N (coding positions 1-723) and pGEX6P1-SF3B1-C (724-3300) by amplifying the coding sequences from *T. brucei* genomic DNA and inserting them into the EcoRI/XhoI and BamHI/XhoI restriction sites of the pGEX6P1 vector (Cytivia), respectively. To mutate the 13 threonine codons of the TP motifs to alanine codons in the N-terminal domain (NTD) of SF3B1 (Figure S1), we had the mutated NTD sequence synthesized (IDT). Amplification products of this APmut-termed sequence were transferred into the expression vectors to obtain pT7-SF3B1-APmut-HA-BLA and pGEX6P1-SF3B1-N-APmut. Maps of newly constructed plasmids are provided in Figure S2.

### Trypanosomes

Insect-stage, procyclic *T. brucei brucei* Lister strain 427 was grown in SDM-79 medium supplemented with 10% fetal bovine serum as described previously (41). For stable transfections, 10^8^ cells were mixed with 10 μg of linearized plasmid, electroporated, and immediately cloned by limiting dilution as detailed before (42). PCR of genomic DNA with one primer annealing outside the cloned region was used to confirm correct plasmid integration. For selection, antibiotics G418, hygromycin, phleomycin and blasticidin were added to the medium at final concentrations of 40, 40, 2.5 and 20 μg/ml, respectively. In 29-13 trypanosomes and its derivative cell lines, the respective concentrations of G418 and hygromycin were 15 and 50 μg/ml. In culture growth assays, trypanosomes were counted manually with a hemocytometer and diluted daily to 2 × 10^6^ cells/ml. Cumulative growth was calculated by extrapolating daily growth rates. To determine the half maximal effective concentration (EC_50_) of OTS964 (Selleck Chemicals) on cell viability, we incubated 10^5^ cells in a 100 μl volume in 96 wells with the inhibitor for 48 h in 24 replicates. The concentration of OTS964 was serially diluted 1:2 from 250 to 0.12 μM, and the number of viable cells indirectly measured using the luminescent CellTiter-Glo^®^ assay (Promega) according to the manufacturer’s specifications. Lowest and highest average chemiluminescence signals were set to 0 and 100, respectively, and the EC_50_ value calculated by non-linear regression using the GraphPad Prism 10 program (data are listed in Table S1).

### Protein analysis

For preparation of whole cell lysates, 1.5 × 10^7^ trypanosomes were pelleted at 2,700 g for 1 min at room temperature and dissolved in 50 μl of standard SDS protein loading buffer. Trypanosome extracts were prepared as described previously (43). To detect PTP- and HA-tagged proteins and α tubulin in immunoblots, the ProtA-specific Peroxidase Anti-Peroxidase (PAP) reagent or the monoclonal HPC4 anti-ProtC antibody, a monoclonal rat anti-HA antibody and a mouse monoclonal anti-[human] α tubulin antibody (all from Millipore Sigma) were used, respectively. CRK9AP was visualized by a previously established polyclonal immune serum from rat (34). Except when PAP reagent was used, peroxidase-conjugated secondary antibodies were applied and blots developed using the BM chemiluminescence blotting substrate (Millipore Sigma) according to the manufacturer’s specifications. GST-tagged proteins were detected with a Dylight 800-conjugated goat anti-GST antibody (Rockland), and signals recorded with a ChemiDoc Imaging System (BioRad).

For the detection of the elusive phosphorylated form of HA-tagged SF3B1, pelleted trypanosomes were resuspended in ice-cold KLB solution (25 mM Tris-HCl pH 7.4, 150 mM NaCl, 5 mM EDTA, 10% glycerol, 1% Triton X-100), containing phosphatase inhibitors sodium pyrophosphate (10 mM), sodium-Orthovanadate (1 mM), glycerol phosphate (10 mM), sodium fluoride (10 mM) as well as phosphate inhibitor cocktail 2 (Millipore Sigma), and lysed by 5 sonication cycles (30 s on*/*30 s off) at high settings in a Bioruptor sonicator (Diagenode). To optimally distinguish phosphorylated and unphosphorylated SF3B1, the samples were separated on NuPAGE 3-8% Tris-acetate gels (Thermo Fisher Scientific) and subjected to immunoblotting. To establish that the slow migrating SF3B1 band in SDS-PAGE corresponded to phosphorylated SF3B1, lysates were gel-filtrated through 7K Zeba Spin Desalting columns (Thermo Fisher Scientific), which were equilibrated in KLB solution with or without phosphatase inhibitors, incubated at room temperature for 20 min, and then mixed with SDS loading buffer and immediately incubated at 100 °C for 5 min to block endogenous phosphatases.

For *in vitro kinase* assays, CRK9**^WT^**-PTP and CRK9**^AS^**-PTP were tandem affinity-purified according to the published protocol (38). As substrates, recombinant GST-SF3B1-N, GST-SF3B1-N-APmut and GST-SF3B1-C were expressed in the *Escherichia coli* DH5α strain and purified with glutathione agarose beads (see Figure S3 for details). Kinase reactions were carried out for 30 min at 28 °C in a volume of 25 µl in kinase buffer (20 mM Tris-HCl pH 7.7, 30 mM KCl, 4 % sucrose, 7 mM MgCl2, 0.1 mg/ml BSA) containing 1 mM ATPγS or 6-cHe-ATPγS (N⁶-cyclohexyladenosine-5’-O-(3-thiotriphosphate)) and 5 µl of final CRK9**^WT^**-P **^or^** CRK9**^AS^**-P eluate. Kinase reactions were stopped by adding EDTA to a final concentration of 2 mM. In a subsequent step, thiophosphates were alkylated by adding 1.5 µl of 50 mM PNBM (p-nitrobenyl mesylate) (Abcam) to the reaction and incubating samples for 2 h at room temperature. Samples were analyzed by immunoblotting with the monoclonal anti-thiophosphate ester antibody [51-8] (Abcam) at 1:2,000 dilution.

For co-immunoprecipitation (co-IP) of CRK9-PTP, 100 μl of extract was diluted with 150 μl of E-buffer (150 mM sucrose, 20 mM HEPES–KOH pH 7.7, 20 mM potassium L-glutamate, 3 mM MgCl_2_, 1 mM dithiothreitol [DTT], 10 μg/ml leupeptin, 10 μg/ml aprotinin), mixed with 30 μl of ProtA-binding, E buffer-equilibrated IgG Sepharose 6 Fast Flow beads (Cytiva), and rotated for 2 hours at 4 °C. Subsequently, beads were washed six times with 800 μl of TT150 buffer (20 mM Tris–HCl pH 8.0, 150 mM NaCl, 3 mM MgCl_2_, 0.1% Tween20), and proteins released from beads with SDS loading buffer.

To determine the OTS964 half maximal inhibitory concentration (IC_50_) of CRK9 autophosphorylation *in vitro*, we carried out 3 independent series of kinase experiments in which we lowered the OTS964 concentration in ten 1:2 dilution steps from 500 to 0.49 μM, analyzed thiophosphorylation by immunoblotting, and quantified signals by densitometry using Image Lab Software Version 6.1 (BioRad). For each series, the signal strength of a reaction without inhibitor was set to 100, and the IC_50_ value calculated by non-linear regression using GraphPad Prism 10 (data are listed in Table S2).

### RNA analysis

To detect *PAP1* pre-mRNA splicing defects, 10^8^ cells were treated with DMSO or inhibitors and total RNA prepared using the Trizol reagent (ThermoFisher Scientific) according to the manufacturer’s protocol. cDNA was generated from 1 μg of RNA by reverse-transcription with SuperScript IV (ThermoFisher Scientific) and random hexamers, and cleaned from RNA with 5 units of RNase H for 20 min. at 37 °C. cDNA was amplified by 2- and 3-primer PCR assays as published (32) to reveal distinct products of spliced and unspliced pre-mRNA for *cis* and *trans* splicing, respectively. DNA oligonucleotides that were used in assays are listed in Table S3.

For RNA immunoprecipitation (RIP) assays, 200 ml cultures of trypanosomes, which expressed PRP19-PTP and analog-sensitive CRK9^AS^-HA and no wild-type CRK9, were grown to ~1 × 10^7^ cells/ml and treated with DMSO or 10 μM of 1-NM-PP1 for 3 hours. Following extract preparation, PRP19-PTP was precipitated from 100 μl of extract that was diluted with 400 μl of PA150 buffer (150 mM KCl, 20 mM Tris-HCl pH 7.7, 3 mM MgCl_2_, 0.5 mM DTT, 0.1% Tween20) and combined with 30 μl of pelleted, PA150-equilibrated, ProtA-binding IgG Sepharose 6 Fast Flow beads (Cytiva), by rotating the reaction for 1 hour at 4 °C. After washing the beads five times with 800 μl of PA150 buffer, total RNA was prepared from pellets using the Trizol reagent. For primer extension assays of SL RNA and U2 snRNA, 3 μg of input RNA prepared from extract or a third of the precipitated RNA was annealed to 5 ng of 5’-biotin labeled DNA oligonucleotides bio-SL_PE and bio-U2_PE, and reverse-transcribed with Superscript IV. Primer extension products were separated on a 50% urea/6% polyacrylamide gel, transferred to a positively charged nylon membrane (Roche) by electroblotting, and detected with the Chemiluminescent Nucleic Acid Detection Module Kit (ThermoFisher) according to the manufacturer’s protocol.

For quantitative analyses of U snRNA and mRNA abundances, total RNA was reverse-transcribed with random hexamers and oligo(dT)_20_, respectively, and cDNA subjected to qPCR. qPCR reactions were performed with SsoFast EvaGreen Supermix (Biorad) on a CFX96 cycler (Biorad). Oligonucleotide pairs were considered suitable for qPCR when they produced a single product of correct size in agarose gel electrophoresis, exhibited a single peak in melting curve analysis, and revealed a coefficient of determination (*R*^2^) in a standard curve of serially diluted input samples between 0.98 and 1.0. To determine RIP efficiency, three independent RIP assays were conducted with parallel DMSO and 1-NM-PP1 treatments. U snRNA abundances in precipitates were corrected by those determined for inputs, and, in each case, the precipitates were directly compared and the DMSO-derived value set to 100. The *SF3B1* knockdown efficiency in gene silencing experiments was determined one day after dox treatment, comparing S*F3B1* mRNA abundance to that of untreated cells and normalizing values with those of α tubulin mRNA.

### Protein structure prediction

Structural models of the CRK9-OTS964 complexes were generated using Chai-1 (44). The SMILES representation of OTS964 was prepared with Open Babel (45) using the PDB coordinates from the structure of the CDK11B-OTS964 complex (PDB ID: 7UKZ). For each complex, five models were predicted and the top-ranked model was selected for analysis and figure preparation. The accuracy of the predicted complexes was assessed using pTM (predicted template modeling) and ipTM (interface template modeling) scores. Structural alignment and molecular visualization were performed using PyMOL (Schrödinger, Inc.). The volumes of the inner cavities of the complexes were calculated with CASTp (46) in the absence of OTS964, using a cavity detection probe radius of 1.8 Å.

## Results

### CRK9 activity is essential to maintain SF3B1 phosphorylation *in vivo*

To study SF3B1 phosphorylation in trypanosomes, we employed analog-sensitive kinase technology (47,48). Briefly, kinases have a large amino acid residue in their ATP-binding pocket known as the gatekeeper. When this residue is replaced by a smaller amino acid such as glycine, the extra space typically renders the kinase sensitive to derivatives of the small molecule kinase inhibitor PP1 and enables the analog-sensitive kinase to effectively use bulky ATP analogs, which are too large for wild-type kinases, as substrates.

As we demonstrated previously, the gatekeeper mutation M438G rendered CRK9 analog sensitive. By establishing the procyclic cell lines TbCRK9**^WT^**-PTPee and TbCRK9**^AS^**-PTPee, which exclusively express wild-type and analog-sensitive CRK9 (CRK9**^AS^**) with a C-terminal PTP tag, we showed that adding 1-NM-PP1 to medium specifically inhibited CRK9**^AS^**-PTP, leading to a rapid defect in both *trans* and *cis* splicing (32). To enable detection of SF3B1 in these two cell lines, we stably transfected them with the linearized plasmid PURO-HA-SF3B1, targeting it for integration into an endogenous *SF3B1* allele (Figure 1A). Accordingly, the resulting two cell lines expressed HA-SF3B1 in two distinct bands, a stronger band of correct size (~123 kDa) and a weaker band of larger size. As expected, both cell lines maintained the expression of CRK9-PTP (Figure 1B). To demonstrate that the upper band represents phosphorylated SF3B1, we first prepared extract and found that the band rapidly disappeared, an observation that is consistent with experiments in the human system (9). To circumvent this, we used whole cell lysates, removed detergent by gel filtration and showed that stabilization of the upper band required phosphatase inhibitors which indicated that the upper band does represent phosphorylated HA-SF3B1 (Figure 1C).

**Figure 1.**
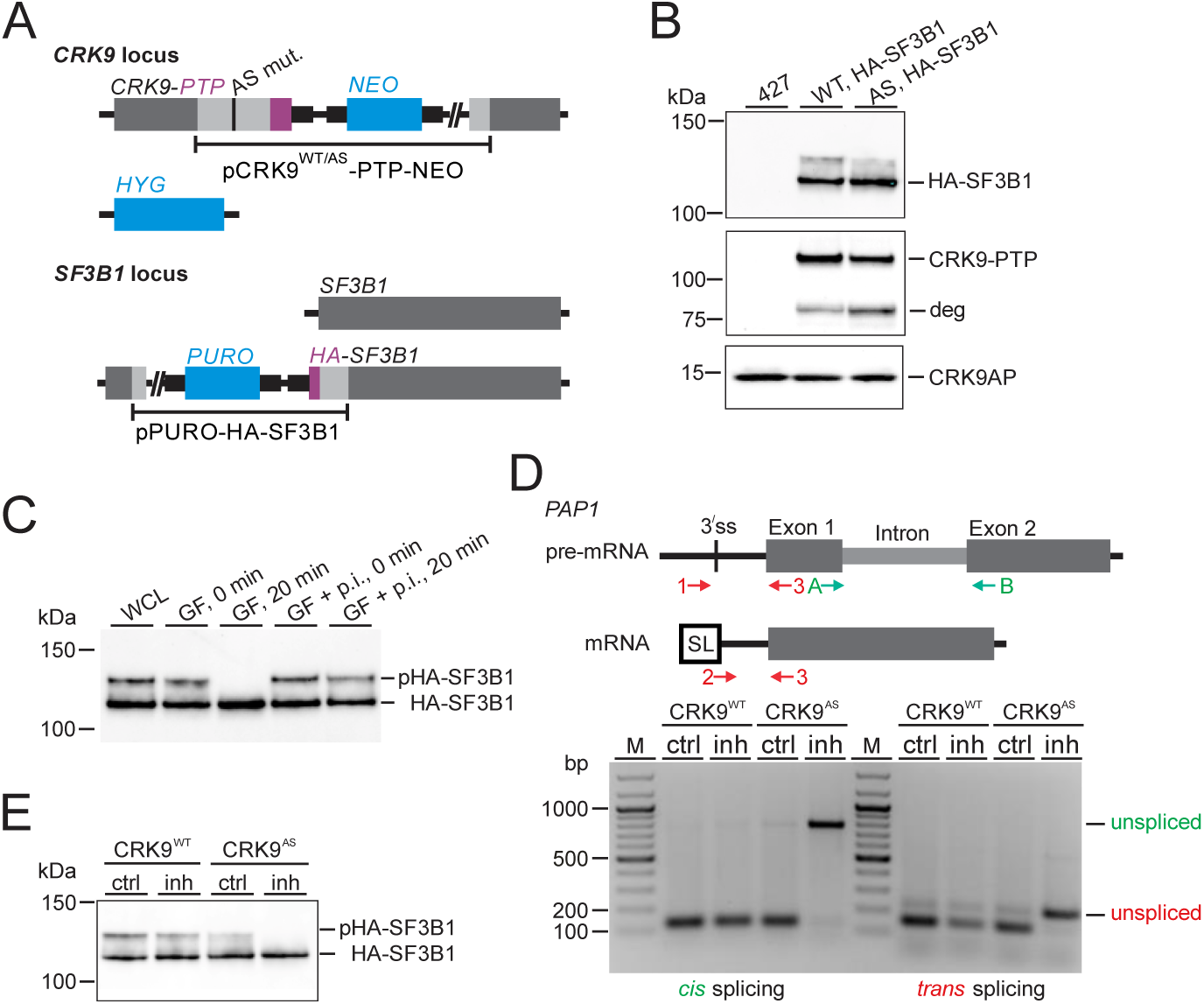
Trypanosome SF3B1 is phosphorylated in CRK9-dependent manner. (**A**) Schematic depiction of the modified CRK9 and SF3B1 loci in the CRK9^WT/AS^ cell lines expressing HA-SF3B1. Plasmids CRK9^WT^-PTP-NEO or CRK9^AS^-PTP-NEO were integrated into one *CRK9* allele and the remaining wild-type allele was replaced by the hygromycin phosphotransferase gene (*HYG*). In each cell line, integration of pPURO-HA-SF3B1 into one *SF3B1* allele fused the HA tag N-terminally to the splicing protein. (**B**) Immunoblot showing the expression of HA-SF3B1 (top panel, anti-HA antibody), CRK9-PTP (middle panel, PAP reagent detecting the tag), and CRK9AP (anti-CRK9AP immune serum) in unmodified cells (427) and in CRK9^WT^-PTP (WT) and CRK9^AS^-PTP (AS) expressing cell lines. deg, putative CRK9-PTP degradation product. (**C**) Anti-HA immunoblot of whole cell lysate (WCL) that was subjected to gel filtration (GF) in the absence or presence of phosphatase inhibitors (p.i.) and incubated for 0 or 20 minutes. The top band corresponds to phosphorylated (p)HA-SF3B1. (**D**) 2- and 3-primer PCR assays to monitor *cis* and *trans* splicing defects of *PAP1* RNA in WT and AS cells that were treated with DMSO or 10 μM 1-NM-PP1 (inh) for 3 hours. (**E**), Anti-HA immunoblot of WT and AS whole cell lysates after treatment with DMSO (ctrl) or 1-NM-PP1 (inh).

To confirm that the new cell lines maintained their differential susceptibility to the inhibitor, we treated the cells with 10 μM of 1-NM-PP1 for three hours and employed established 2- and 3-primer RT-PCR assays to analyze *cis* and *trans* splicing defects specifically of the intron-containing *PAP1* pre-mRNA. As anticipated, 1-NM-PP1 strongly affected both splicing events only in cells that expressed CRK9**^AS^**-PTP but not in those expressing CRK9**^WT^**-PTP (Figure 1D). Finally, immunoblotting of cell lysates revealed that blocking the kinase activity of CRK9**^AS^**-PTP led to a loss of phosphorylated HA-SF3B1 (Figure 1E). We therefore concluded that CRK9 activity is required for SF3B1 phosphorylation.

### CRK9 directly phosphorylates TP motifs in the N-terminus of SF3B1

Human SF3B1 has an intrinsically unstructured N-terminal domain (NTD) that spans amino acids 1 to 463 and contains 29 TP motifs, followed by a long HEAT repeat domain and a C-terminal anchor region (8). *In vitro* kinase assays showed that SF3B1 phosphorylation by human CDK11 is restricted to the NTD, and CDK11 inhibition *in vivo* reduced the phosphorylation of four TP motifs for which specific antibodies were available (13). A sequence alignment of human, *T. brucei* and *Leishmania major* SF3B1 indicated that the trypanosome N-terminal domain spans amino acids 1 to 241 and contains 13 TP motifs. The large HEAT repeat region extends from amino acids 307 to 984 and is moderately conserved in sequence, whereas the anchor region, comprising amino acids 1042-1099, is highly conserved (Figure S4).

To determine whether CRK9 directly phosphorylates the NTD of SF3B1 we developed an *in vitro* kinase assay with tandem affinity-purified CRK9**^WT^**-P or CRK9**^AS^**-P (please note that the PTP tag was reduced to P after TEV protease cleavage during tandem affinity purification) and substrates GST-SF3B1-N (aa 1-241) and GST-SF3B1-C (aa 242-1099, Figure 2A), which were expressed in and purified from *E. coli* (Figure S3). To demonstrate CRK9-dependent phosphorylation, we performed thiophosphorylation reactions using either ATPγS or its bulkier analog 6-cHe-ATPγS which can be efficiently used by analog-sensitive but not wild-type CRK9 (Figure 2B). After the reactions, thiophosphates were alkylated with PNBM and detected on immunoblots with an anti-thiophosphoester antibody. Please note that CRK9 is an autophosphorylating kinase, adding a phosphate likely to the T533 residue in its activation loop (31,34,49). As shown in Figure 2B, kinase reactions with ATPγS enabled both wild-type and analog-sensitive CRK9 to autophosphorylate, and both enzymes phosphorylated GST-SF3B1-N but not GST-SF3B1-C or GST alone (lanes 5-10). The lower efficiency of CRK9**^AS^**-P in utilizing ATPγS is most likely due the introduced extra space in its ATP-binding pocket and consistent with previous results (32). On the other hand, 6-cHe-ATPγS proved to be too bulky for wild-type CRK9 because in the presence of this nucleotide, auto and GST-SF3B1-N phosphorylation was nearly abolished (lanes 11,13, 15). In contrast, CRK9**^AS^**-P was able to efficiently use the bulky ATP analog in phosphorylating itself and GST-SF3B1-N (lanes 12,14, 16). Taken together, these results demonstrated that CRK9 specifically phosphorylated the NTD of SF3B1 and not the remainder of the protein. Importantly, the dependence of GST-SF3B1-N thiophosphorylation on CRK9**^AS^**-P when 6-cHe-ATPγS was used excludes the possibility that phosphorylation was carried out by another endogenous kinase that was co-purified with CRK9.

**Figure 2.**
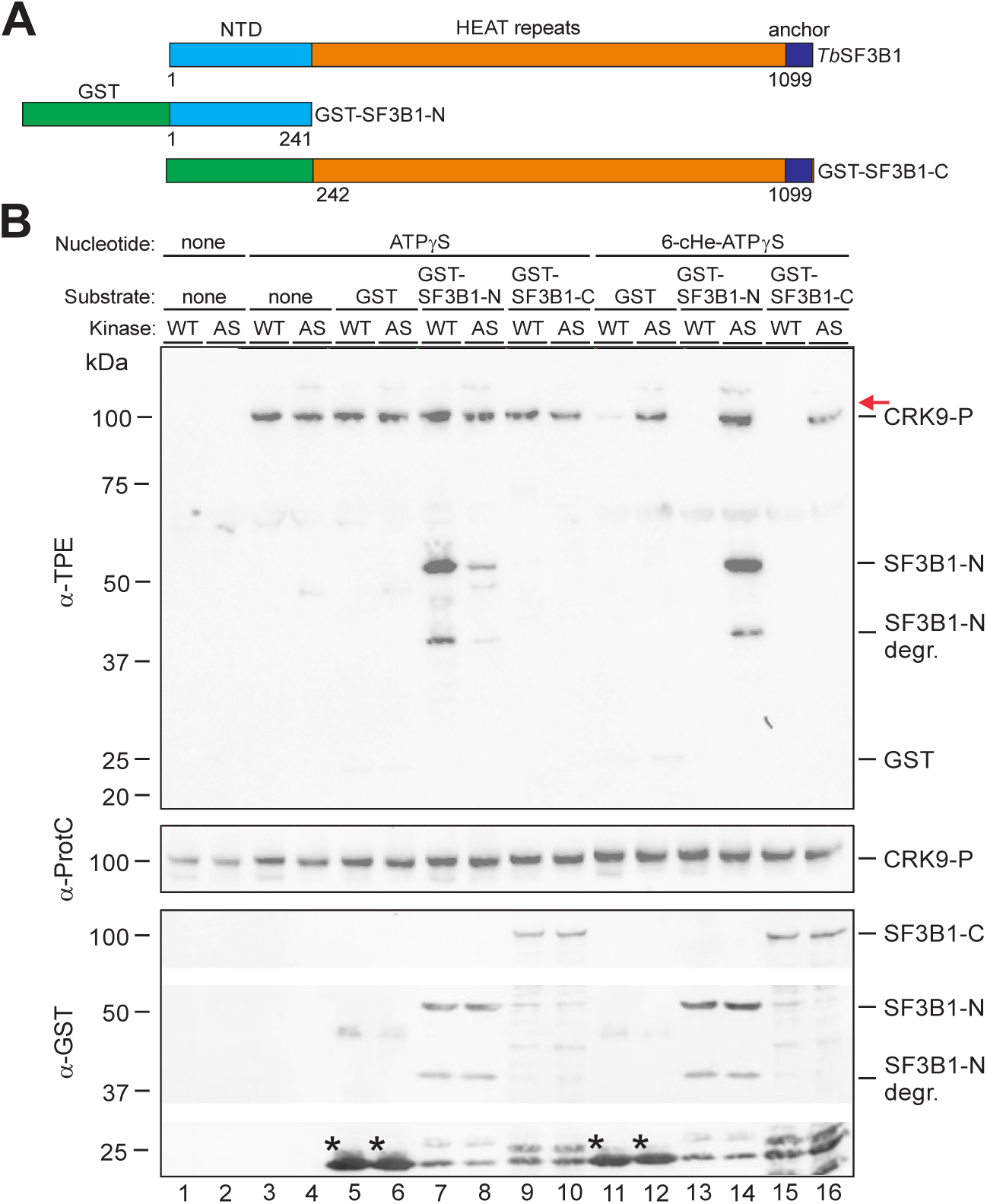
CRK9 directly phosphorylates the NTD of SF3B1 *in vitro*. (**A**) Schematic of *T. brucei* SF3B1 and the fusion proteins GST-SF3B1-N and GST-SF3B1-C with N-terminal domain (NTD), HEAT repeat domain, anchor region and GST tag depicted to scale. (**B**) *In vitro* kinase assays were carried out with endogenous, tandem affinity-purified CRK9**^WT^**-P (WT) or CRK9**^AS^**-P (AS) enzyme, substrates GST, GST-SF3B1-N or GST-SF3B1-C in the presence of ATPγS or its bulky analog 6-cHe-ATPγS. Kinase reactions were analyzed by immunoblotting. Thiophosphorylated proteins after alkylation with PNBM were detected with anti-thiophosphate ester antibody 51-8 (α-TPE), CRK9-P with anti-ProtC antibody, and GST fusion proteins with anti-GST antibody (α-GST). A degraded (degr.) GST-SF3B1 product of ~40 KDa was consistently detected in our samples. In the α-TPE blot CRK9-P is detected due to autophosphorylation and the red arrow indicates the expected position of GST-SF3B1-C. Asterisks identify GST control substrate.

Next, we wanted to determine which residues in the NTD were phosphorylated by CRK9. The NTD of human SF3B1 is heavily phosphorylated. Specifically, 32 threonines, 26 of which reside in TP motifs, plus 15 serine and 2 tyrosine residues have been identified as phospho-sites (50–54). The NTD of trypanosome SF3B1 harbors, in addition to the 13 TP motifs, 18 serine, an additional 12 threonine and six tyrosine residues as potential phosphorylation sites. However, unambiguous identification of specific phosphopeptides in our *in vitro* assays by mass spectrometry has been challenging, likely because only a small fraction of the substrate was phosphorylated. We therefore adopted a different approach. Cyclin-dependent kinases are proline-guided serine/threonine kinases, preferentially phosphorylating TP and SP motifs. Since the NTD in *T. brucei* does not harbor an SP motif, we hypothesized that CRK9 phosphorylation of SF3B1 is restricted to TP motifs. To test this, we expressed and purified GST-SF3B1-N-APmut, in which the 13 TP motifs were mutated to alanine-proline (AP) motifs. In contrast to wild-type GST-SF3B1-N, we could not detect any phosphorylation of the mutated substrate (Figure 3A). Since the SF3B1 NTD is intrinsically unfolded, it is unlikely that a folding issue prevented phosphorylation of the mutated substrate. Hence, this result strongly indicates that CRK9 specifically phosphorylates TP motifs in the SF3B1 NTD. Large scale phosphoproteomics support this view because the three phospho-sites identified in trypanosome SF3B1 so far - T61, T63 and T74 - reside in N-terminal TP motifs (55).

**Figure 3.**
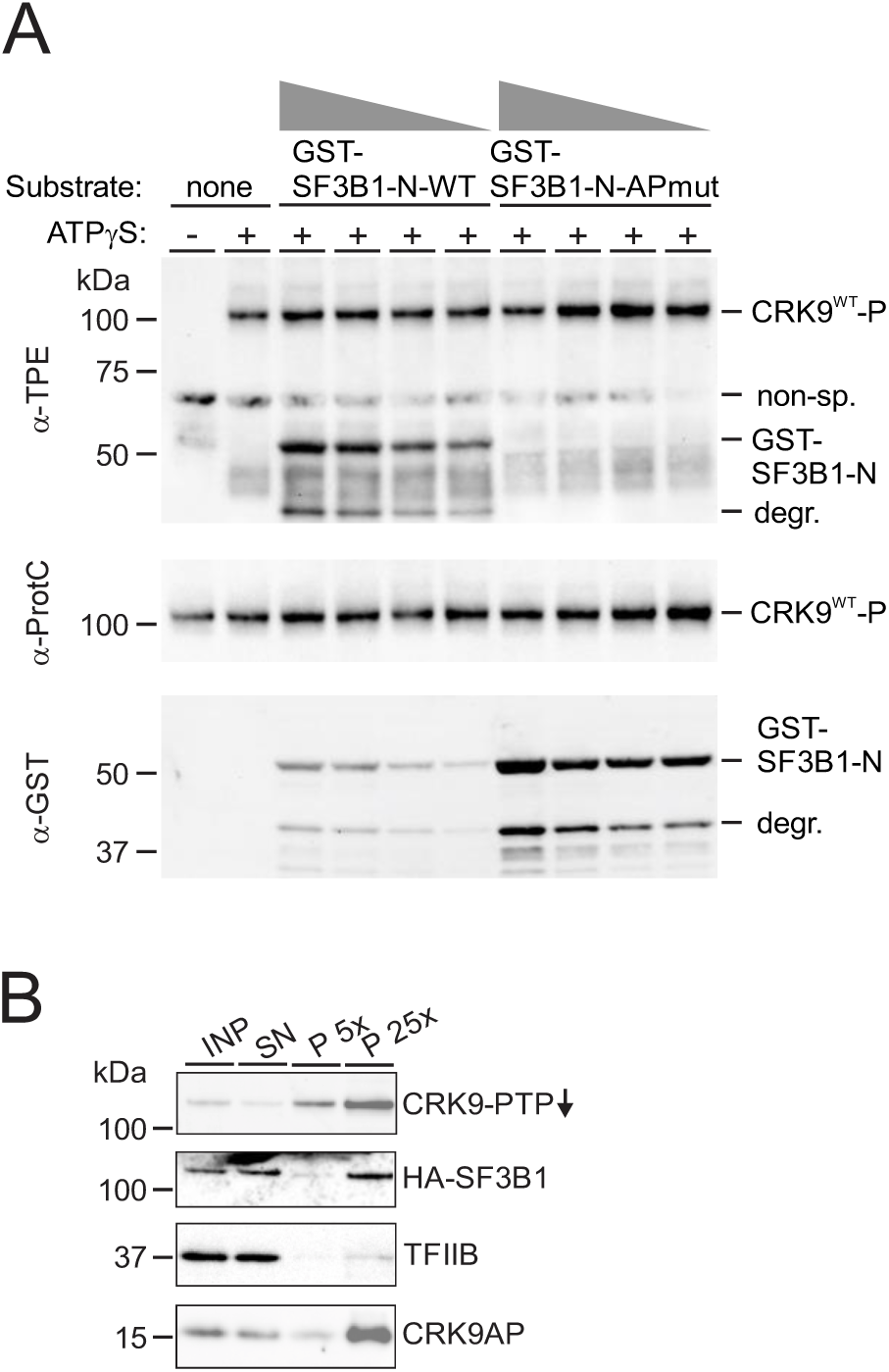
CRK9 phosphorylates TP motifs in the NTD of SF3B1 and interacts with SF3B1 *in vivo*. (**A**) *In vitro* kinase reactions were carried out with endogenous, tandem affinity-purified CRK9**^WT^**-P and decreasing concentrations of wild-type GST-SF3B1-N(-WT) or of GST-SF3B1-N-APmut. A non-specific band (non-sp.) was detected with a-TPE antibody even when ATPγS was omitted in a control reaction. (**B**) CRK9-PTP was immunoprecipitated and detected in immunoblots in extract (INP) supernatant (SN) and precipitate (P) with the PAP reagent. Co-precipitated HA-SF3B1 was detected with an anti-HA antibody, and TFIIB and CRK9AP were detected as negative and positive controls, respectively, with previously raised and characterized polyclonal immune sera. x-Values denote relative amounts analyzed.

If CRK9 directly phosphorylates SF3B1, enzyme and substrate need to physically interact. However, SF3B1, other subunits of the SF3B complex (56,57) or components of the U2 snRNP did not detectably co-purify with CRK9 in previous tandem affinity purifications, indicating that the interaction is highly transient, not very stable and/or occurs only with a small fraction of the substrate (34). We therefore employed co-IP using extract of cells that express CRK9**^WT^**-PTP and HA-SF3B1, a method which can be conducted more rapidly than TAP (Figure 3B). Although it required a relatively large amount of pellet to detect the interaction, the CRK9 pull-down co-precipitated 6.75-fold more of HA-SF3B1 than of TFIIB, the negative control, relative to the input signal from extract (average of two experiments). The rather low efficiency of SF3B1 co-IP is most likely explained by the fact that SF3B1 phosphorylation occurs at a specific stage of spliceosome assembly, apparently in the B complex before the spliceosome is activated (9,13). Hence, access of the kinase to SF3B1 may be restricted to this spliceosomal stage, reducing co-IP efficiency. Together, the *in vitro* kinase reactions and the detectable interaction of CRK9 and SF3B1 in extract strongly support the notion that trypanosome CRK9 and human CDK11 are functional homologs in SF3B1 phosphorylation.

### Inhibition of CRK9 kinase activity interferes with spliceosome activation

In the human system, inhibition of CDK11 prevented spliceosome activation, specifically the formation of the B^act^ complex (13). In trypanosomes, however, characterization of specific spliceosome complexes has not been achieved yet. Nonetheless, RNA immunoprecipitation (RIP) of the spliceosomal protein PRP19 showed that it predominantly associated with the active spliceosome in trypanosomes because it efficiently co-precipitated U2, U5, and U6 snRNA but not U4 snRNA (26,37). Thus, if formation of the active spliceosome is dependent on CRK9 kinase activity, CRK9 inhibition should reduce the interaction of PRP19 with the spliceosome, i.e. these snRNAs. To analyze this, we generated a cell line that expressed PRP19-PTP from one endogenous allele and exclusively expressed CRK9^AS^-HA (Figure 4A). We treated these cells either with DMSO or with 10 μM 1-NM-PP1 for 3 hours, prepared extract, and carried out anti-PRP19-PTP RIP assays. Despite the short treatment period, a primer extension assay showed a substantial increase of full-length SL RNA and a concomitant decrease of the SL intron/Y structure intermediate signal in extract while the signals for U2 snRNA were of comparable strength (Figure 4B). This result supports our previous observation obtained with *CRK9* silencing that the kinase activity is fundamentally important for the first step of pre-mRNA splicing (31). As anticipated, the pull-down of PRP19-PTP co-precipitated less U2 snRNA after CRK9 inhibition. To quantify the RIP efficiencies of U2, U5 and U6 snRNAs, we carried out four independent RIP assays and quantified precipitated U2 snRNA relative to input by RT-qPCR. As shown in Figure 4C, PRP19 association with those U snRNAs was significantly reduced when CRK9^AS^ activity was blocked by 1-NM-PP1. This finding indicates that CRK9 kinase activity is required for spliceosome activation in trypanosomes.

**Figure 4.**
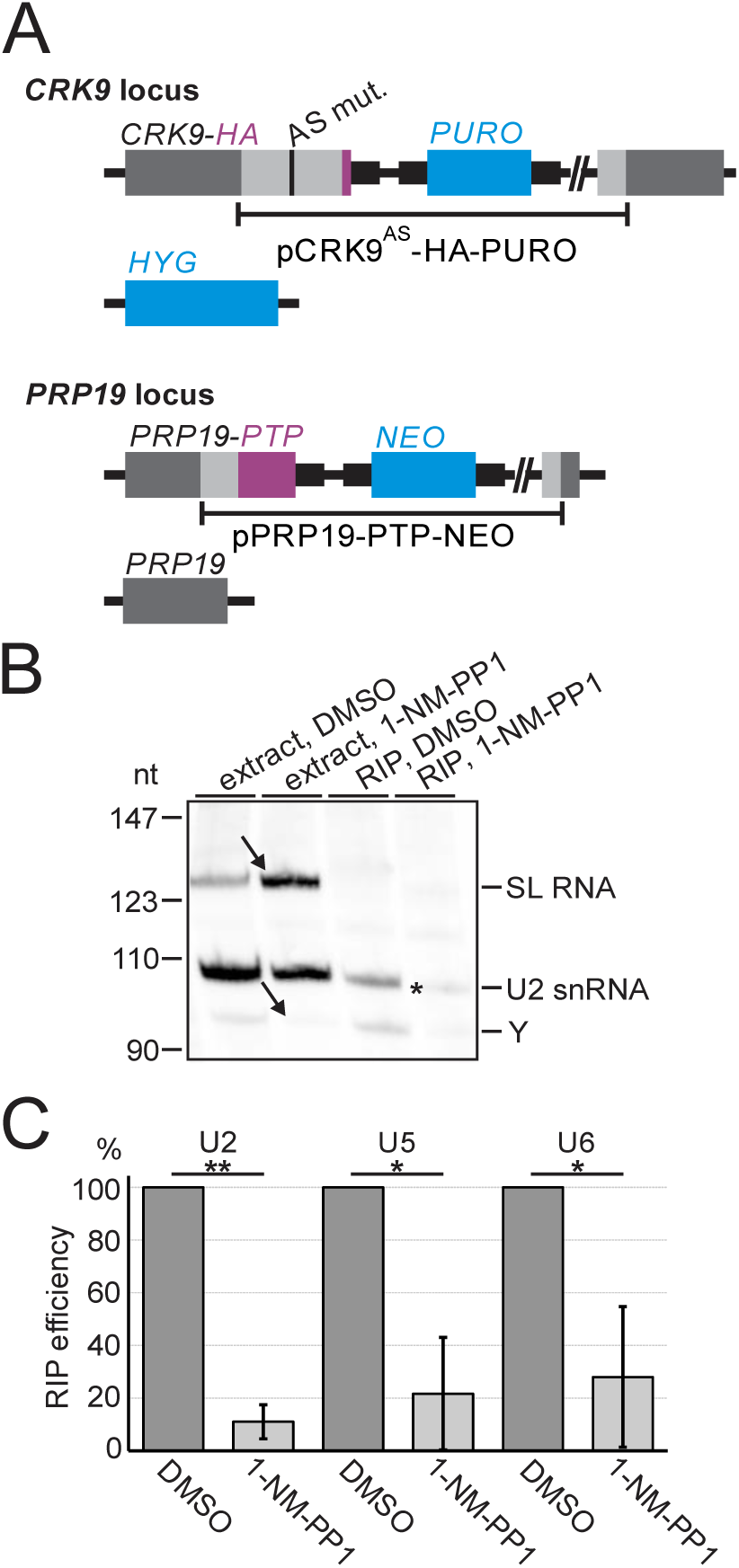
Inhibition of CRK9^AS^ with 1-NM-PP1 reduces the interaction of PRP19 with the active spliceosome. (**A**) Schematic depiction of the CRK9 and PRP19 loci in the procyclic trypanosome line CRK9^AS^-HAee/PRP19-PTP which exclusively expresses analog-sensitive, HA-tagged CRK9 and, from one endogenous allele, C-terminally PTP-tagged PRP19 (PRP19-PTP). (**B**) Cells were incubated for 3 hours in the presence of 10 μM 1-NM-PP1 or DMSO and subsequently used to prepare extracts. In RNA immunoprecipitation (RIP) assays, PRP19-PTP was precipitated with IgG beads, and total RNA preparations subjected to primer extension assay with biotinylated oligonucleotides SL_PE and U2f which are antisense to SL RNA and U2 snRNA, respectively. Primer extension products were separated on an 8% polyacrylamide/50% urea gel, blotted and detected with a streptavidin-peroxidase conjugate. Y denotes the SL *trans* splicing intermediate. Arrows depict the increased full-length SL RNA product and the decreased Y product. (**C**) RT-qPCR assays of total RNA prepared from extract RIP precipitates, amplifying U2, U5, and U6 sequences.

### The TP motifs in the NTD of SF3B1 are essential for pre-mRNA splicing

Since our data suggested that CRK9 phosphorylation is restricted to TP motifs in the SF3B1 NTD, we wanted to know whether these motifs are important for pre-mRNA splicing. For the corresponding analysis we first generated a procyclic cell line for conditional *SF3B1* silencing through the RNA interference (RNAi) pathway. We stably transfected so-called 29-13 cells, which constitutively express the tetracycline repressor and T7 RNA polymerase (58), with plasmid T7-SF3B1-3^/^UTR-stl for dox-controlled expression of a hairpin RNA that targets the 3^/^ UTR of SF3B1 for mRNA degradation. In a second step, we investigated whether the stable transfection of *SF3B1* transgenes, which encoded a C-terminal HA tag and a different, RNAi-resistant 3^/^ UTR could rescue the culture growth and splicing defects. As transgenes we transfected either the wild-type SF3B1 gene (WT) or a mutated SF3B1 gene (APmut) in which the thirteen TP motifs in the NTD were converted to AP motifs (Figure 5A). Since we found it likely that constitutive expression of APmut could stall spliceosome assembly, exerting a dominant negative effect, we placed the transgenes under the control of a dox-inducible T7 promoter. In this way, the presence of dox simultaneously led to silencing of endogenous *SF3B1* and expression of the *SF3B1-HA* transgene.

**Figure 5.**
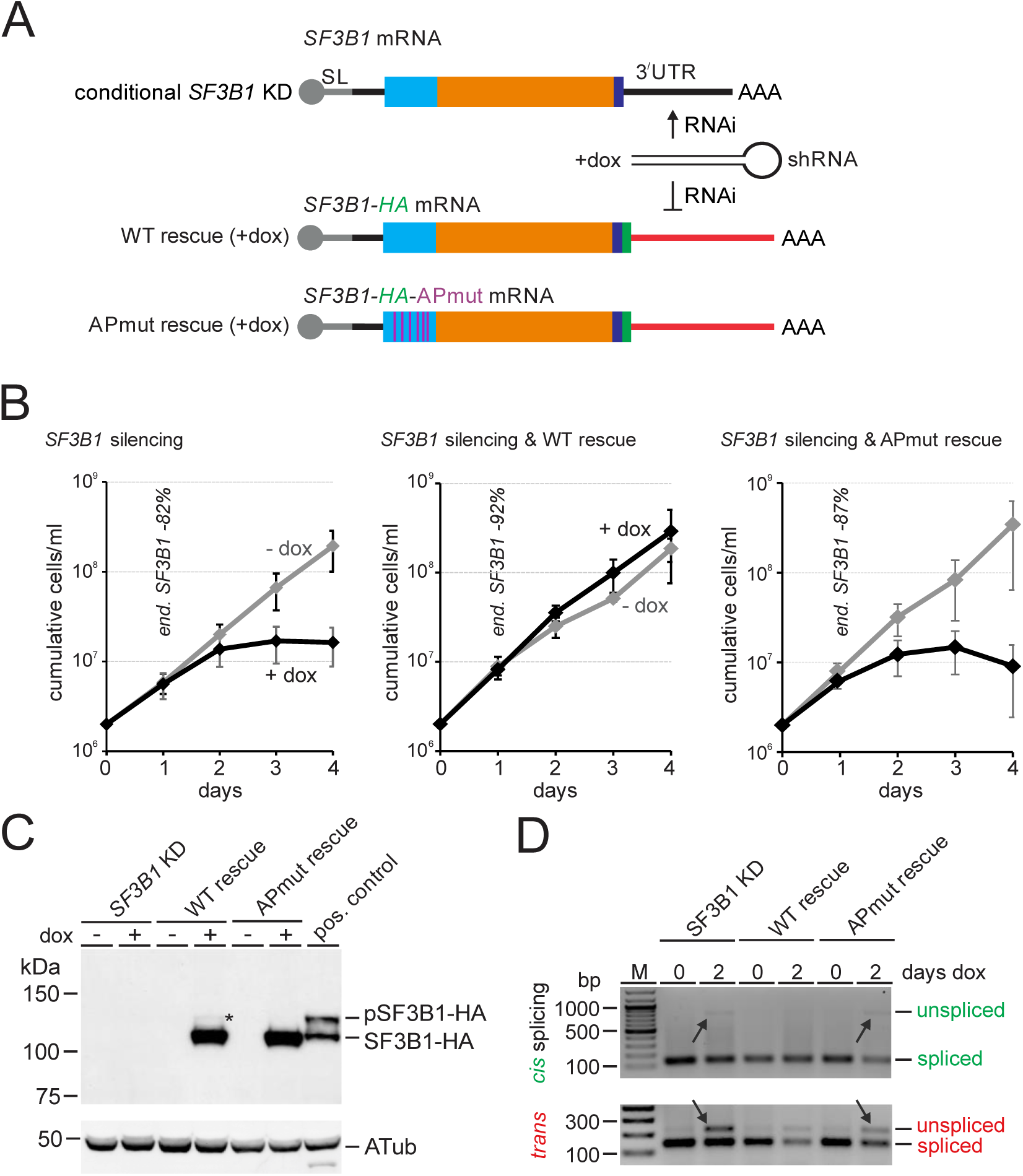
The TP motifs in SF3B1’s NTD are of crucial importance to cell viability and pre-mRNA splicing. (**A**) Schematic outline of the experimental strategy. First, a cell line for conditional *SF3B1* silencing was generated in which dox induces the expression of a hairpin RNA that targets the 3^/^ UTR of endogenous SF3B1 mRNA for degradation by the RNAi pathway. In second steps, complete RNAi-resistant *SF3B1* transgenes under the control of a dox-controlled promoter were stably transfected into this cell line to determine their rescue potential. The transgenes were either wild-type (WT) or mutated such that the 13 TP motifs of the NTD (blue) were changed to AP motifs (APmut); they carried the HA tag sequence at the 3’ end of their coding regions and a different, RNAi-resistant 3^/^ UTR. (**B**) Culture growth of three independently derived clonal lines for each type of cell line in the absence and presence of dox were monitored for four days. The knockdown efficiency was determined by RT-qPCR after 1 day of induction for one representative cell line for each type, using α tubulin mRNA for normalization. (**C**) Immunoblot of whole cell lysates of cell lines treated with or without doxycycline for 1 day, detecting SF3B1-HA, including its phosphorylated form (pSF3B1-HA, asterisk) with a tag-specific antibody and α tubulin as a loading control. (**D**) *PAP1 cis* and *trans* splicing RT-PCR assay of the three cell lines which were untreated or treated with dox for 2 days.

We obtained three independently derived clonal cell lines for each the *SF3B1* knockdown, the WT rescue and the APmut rescue. As anticipated for a general splicing factor, *SF3B1* silencing caused cultures to halt growth after 3 days of dox treatment, a phenotype that was rescued by the expression of wild-type but not of APmut *SF3B1-HA,* indicating the functional importance of the TP motifs (Figure 5B). Immunoblotting showed that the inducible SF3B1-HA transgenes were overexpressed when compared to lysates from cells in which HA-SF3B1 was expressed from an endogenous allele. While this transgene overexpression may have resulted in the relative minor amount of phosphorylated HA-SF3B1 in the wild-type rescue, this band was not detected in the APmut rescue, confirming our *in vitro* kinase results, that CRK9 specifically phosphorylates TP motifs in the NTD (Figure 5C, asterisk). In a final step we wanted to confirm that the growth defects were caused by impaired pre-mRNA splicing, employing the *PAP1* RT-PCR assays. Although the signals of unspliced products in these *SF3B1* knockdown experiments were not as pronounced as those upon chemical inhibition of analog-sensitive CRK9 (see Figures 1D, 6B), which may be due to different kinetics in the longer period until a gene knockdown becomes effective, the impairment of *PAP1 cis* and *trans* splicing was clearly apparent in the *SF3B1* knockdown and in the failed rescue with *SF3B1-HA-APmut* (Figure 5D). These results provide independent evidence for the TP motifs being essential for splicing and for CRK9-mediated phosphorylation of these motifs being required for spliceosome activation.

**Figure 6.**
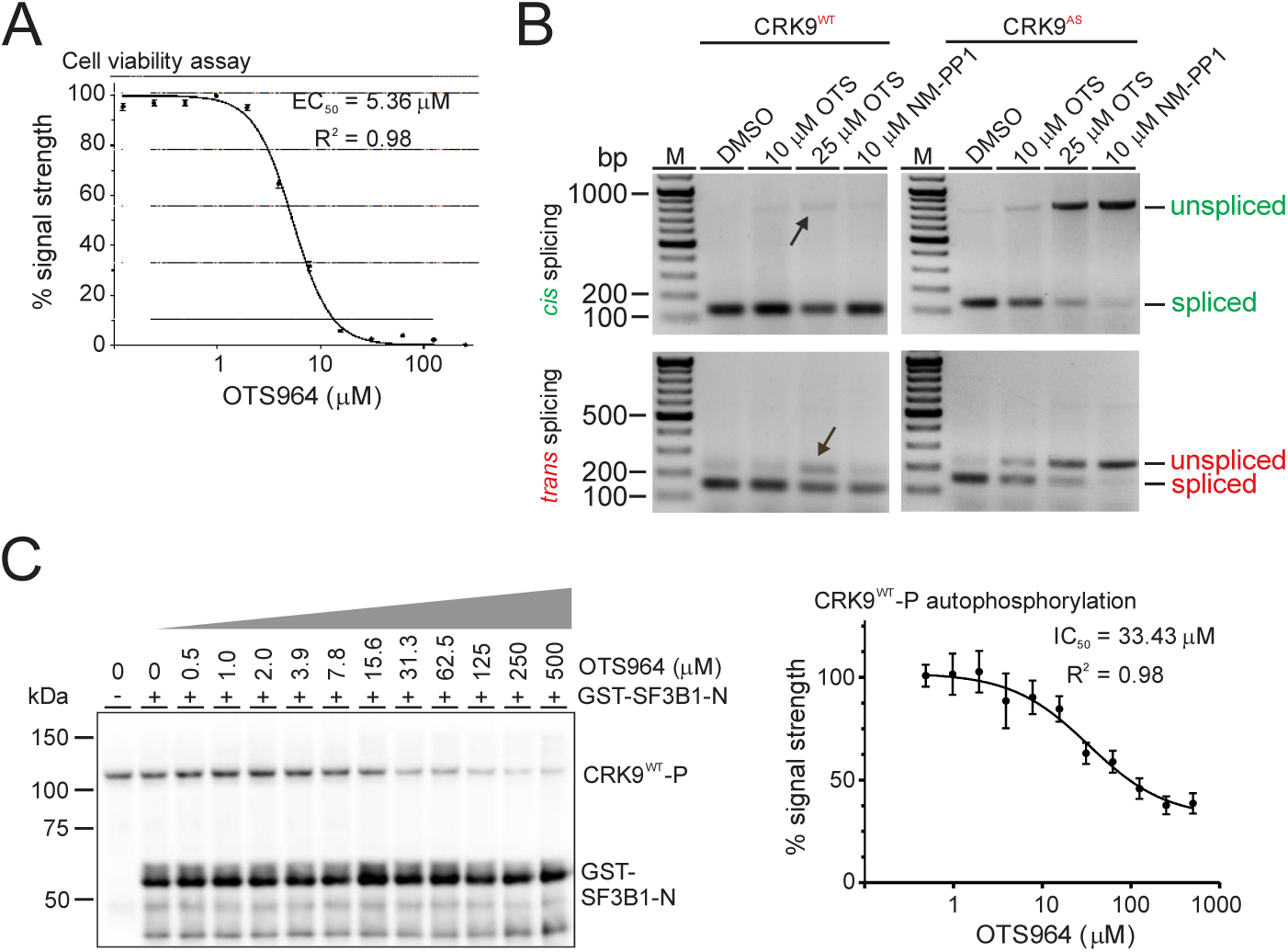
Trypanosomes and CRK9 are insensitive to OTS964. (**A**) Non-linear regression EC_50_ analysis of fluorescence measurements using a viability assay after cells were incubated at different OTS964 concentrations for 48 h. (**B**) Cell lines which exclusively express either CRK9^WT^-PTP or CRK9^AS^-PTP were treated with DMSO, 10 or 25 μM OTS964 or 10 μM 1-NM-PP1 for 3 h. Total RNA preparations of these cells were then reverse-transcribed with random hexamers and subjected to 2- and 3-primer PCR assays to monitor *PAP1* splicing defects. Arrows indicate minor splicing defects in CRK9^WT^-PTP-expressing cells at the highest OTS964 concentration. (**C**) Left, representative immunoblot in which thiophosphorylated CRK9^WT^-P (autophosphorylation) and GST-SF3B1-N were detected after *in vitro* kinase assays with ATPγS and increasing concentrations of OTS964. Right, the IC_50_ of OTS964 on CRK9^WT^-P autophosphorylation was determined by measuring the immunoblot signal strengths of three biological replicates by densitometry. Please note that it required an OTS964 concentration of 125 μM to reproducibly detect a mild reduction of the GST-SF3B1-N signal.

### CRK9 is largely insensitive to OTS964

Thus far, our data demonstrated that, despite the divergence, CRK9 and CDK11 are functional homologs in phosphorylating the SF3B1 NTD as an essential step in pre-mRNA splicing. The divergence, however, raises the possibility that both enzymes are differently susceptible to kinase inhibitors. To test this hypothesis, we analyzed the potency of OTS964 in inhibiting trypanosome culture growth and CRK9 kinase activity *in vitro*. In the human system, OTS964 was tested against various cancer cell lines, in which the half-inhibitory concentration (IC_50_) ranged between 10 and 100 nM, and the IC_50_ of the CDK11 *in vitro* kinase assay was 1.08 nM (12,13). To make sure that our OTS964 batch was similarly effective, we applied the inhibitor to the human HOP-62 non-small cell lung cancer cell line, which is part of the NCI-60 cell collection, and calculated an EC_50_ of 83.39 nM, which is within the range of previous results (Figure S5). In contrast, the EC_50_ for wild-type procyclic cells of the 427 Lister strain was 5.36 μM in 48-hour growth assays (Figure 6A, Table S1). Moreover, it required 25 μM of OTS964 to cause a very mild *PAP1* splicing defect after 3 h of PTPee cell line, suggesting that OTS964 affects another trypanosome kinase more than CRK9 (Figure 6B). Interestingly, the splicing defect was more pronounced in TbCRK9**^AS^**-PTPee trypanosomes, which indicates that the gatekeeper mutation increased OTS964 susceptibility of trypanosomes, a result consistent with the finding that the gatekeeper residue of human CDK11B is in close proximity to bound OTS964 [(59), see below]. For determining the OTS964 IC_50_ of the CRK9 *in vitro* kinase assay, we serially diluted the inhibitor concentration, starting at 500 μM. While we could detect an inhibitory effect of OTS964 on CRK9 autophosphorylation with an IC_50_ of 33.43 μM, GST-SF3B1-N phosphorylation showed only mild reductions at OTS964 concentrations above 125 μM (Figure 6C). The differential effect may be due to the high concentration of the GST-SF3B1-N substrate in the kinase reaction. In sum, trypanosomes and CRK9 are magnitudes less sensitive to OTS964 than human cells and CDK11.

A 2.6 Å crystal structure of the human CDK11B kinase domain bound to OTS964 was recently solved (59). The inhibitor occupies the ATP-binding pocket and interacts with 21 nearby CDK11B residues eight of which differ in *T. brucei* CRK9 (Figure 7A). More specifically, OTS964 forms hydrophobic interactions with I88, V96, A109, V142, M160 (the gatekeeper residue), Y162 and L213, and hydrogen bonds with V96 and D166. These residues are conserved in CRK9 (I275, V283, A304, V340, M438, Y440, L508 and D444). Because the structure of CRK9 has not been determined experimentally, we used Chai-1, a foundation model for biomolecular structure prediction, to generate structural models of CRK9 and CRK9^AS^ in complex with OTS964 (Figure S6). Overall, the predicted models closely resemble the CDK11B-OTS964 complex; the CRK9 kinase domain can be superimposed onto that of CDK11B with a root mean square deviation of 1.8 A for 271 Cα atoms, which includes superimposition of the two gatekeeper methionine residues (Figures 7B-D). Nonetheless, our analysis suggests three, not mutually exclusive, explanations for OTS964’s inefficiency of inhibiting CRK9 activity. Firstly, it was shown that a G223S mutation in CDK11B caused resistance of human cells to OTS964 and interfered with inhibitor binding, likely by introducing steric hindrance (12,59). The corresponding residue in CRK9 is an even bulkier cysteine (C518), which may exacerbate this effect. Secondly, the predicted ATP-binding pocket of CRK9 is distinctly smaller than that of CDK11B (Figure 7E). In contrast, the pocket of CRK9^AS^ appears markedly enlarged, possibly accounting for its seemingly higher OTS964 susceptibility. Finally, the CRK9-specific insertion spanning residues 349 to 430 is predicted to form a lid-like conformation that could further restrict access of OTS964 to the ATP-binding pocket (Figure 7B, arrow). Together, these analyses provide a structural basis for the refractoriness of CRK9 to OTS964 and highlight opportunities for selective inhibition of the trypanosome kinase.

**Figure 7.**
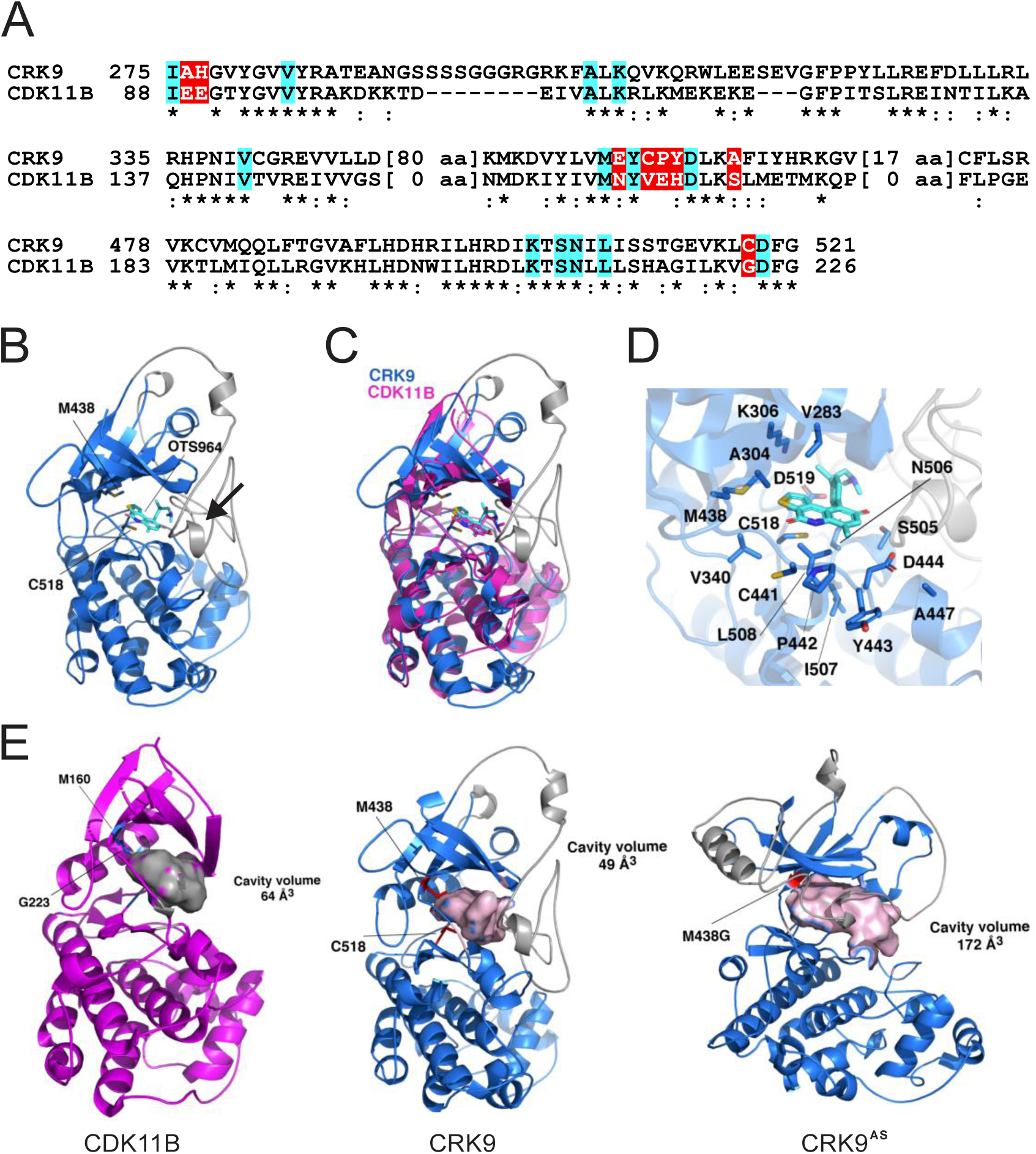
Predicted binding of OTS964 to the CRK9 ATP-binding pocket. (**A**) Sequence alignment around the ATP-binding pocket of the kinase domains of *T. brucei* CRK9 (residues 267-685) and human CDK11B (Uniprot ID: P21127-12; human CDK11B isoform 7). Asterisks indicate identical residues, and colons mark conserved positions. ‘x’ denotes the conserved motif of the cyclin binding helix. The OTS964-interacting residues in CDK11B, identified from the structure of the CDK11B-OTS964 complex (PDB ID: 7UKZ), are highlighted in blue if identical in CRK9 and in red if non-identical. (**B**) Chai-1-predicted model of the CRK9-OTS964 complex. The N-terminal (residues 1-266) and C-terminal (residues 686-744) domains are omitted for clarity. M438, C518, and OTS964 are shown as licorice sticks. The CRK9-specific insertion (residues 349-430) is colored grey. (**C**) Superimposition of CRK9-OTS964 and CDK11B-OTS964 complex structures. CRK9 and CDK11B are shown in blue and magenta, respectively. (**D**) Close-up view of residues surrounding OTS964 in the CRK9 ATP-binding site. (**E**) Molecular surface representation of the OTS964-binding cavities in CDK11B, CRK9, and CRK9^AS^. The cavities were defined using the predicted structures of the CRK9- and CRK9^AS^-OTS964 complexes and the crystal structure of the CDK11B-OTS964 complex. Cavity volumes were calculated using CASTp

## Discussion

Our results demonstrate that spliceosome activation in *T. brucei* depends on CRK9-mediated phosphorylation of TP motifs in the NTD of SF3B1 and that, in this regard, CRK9 is the functional homolog of CDK11. Hence, this essential step for pre-mRNA splicing is of ancient evolutionary origin and not, as has been hypothesized, newly evolved function of metazoan CDK11 (15). It has been argued that the higher complexity of pre-mRNA splicing in metazoans, which have more and larger introns as other eukaryotes, and abundantly carry out alternative splicing, requires additional regulatory proteins such as CDK11. This idea is bolstered by the fact that there are about 80 additional spliceosome-associated proteins in human cells which have no obvious counterpart in *S. cerevisiae* (1). However, it appears that ancestral eukaryotes were intron-rich, potentially having more complex exon-intron structures than extant yeast cells (60). Although *T. brucei* possesses only 3 genes encoding a single intron, in regard to branch point recognition the parasite is complex nonetheless because it harbors roughly ten thousand protein coding genes, all of which are *trans*-spliced and have their own branchpoint upstream of the SL addition site, i.e. the 3^/^ splice site. It therefore appears that the lack of CDK11 in *S. cerevisiae* and its reduced role in *S. pombe* transcription are the result of regressive evolution.

Furthermore, the fact that the kinase activities of CDK11 and CRK9 are globally required for pre-mRNA splicing in human cells and trypanosomes, respectively, SF3B1 NTD phosphorylations arguably reflect a key mechanistic step in spliceosome activation rather than being of regulatory nature. Although we were unable to identify the specific TP sites that were phosphorylated by CRK9, the rather clear separation between phosphorylated and unphosphorylated SF3B1 in gel electrophoresis suggests that more than the three previously identified TP motifs are phosphorylated. Similarly, several TP motifs in the NTD of human SF3B1 were confirmed to be phosphorylated (15). The TP motifs are predominantly spread over the ULM region in human SF3B1 which binds to the core factor U2AF2. U2AF2 is present in trypanosomes (Tb927.10.3500), and it appears that in *T. brucei* SF3B1 at least 3 potential ULMs, including the central tryptophan residue of these motifs, are conserved. Thus, the TP phosphorylations may help to disrupt the interaction between SF3B1 and U2AF2, enabling the release of U2AF2 prior to spliceosome activation. However and in contrast, binding studies with human SF3B1 ULM5 and U2AF2 suggested that TP phosphorylation increased rather than decreased the binding affinity (61). In the human system, the ULM region appears to also interact with the alternative splicing factors CAPERα (RBM39), PUF60 and SPF45 (RBM17) (62,63). Although alternative *trans* splicing has been documented in *T. brucei* (64), no homologs of the latter three factors have been identified, making it unlikely that the fundamentally important splicing step associated with the TP phosphorylations is due to these interactions. Alternatively, it is possible that the discard of U2AF2 during B complex formation may render the SF3B1 NTD accessible for the kinase in the first place. In any case, the massive incoming negative charge deposited by CRK9/CDK11 likely enables SF3B1 to undergo specific interactions through its NTD that are crucially important for spliceosome activation. Unfortunately, and as has been described before (15), the intrinsically disordered nature of the NTD has prevented its resolution in cryo-EM structures of the activated spliceosome. Thus, direct interactions of the NTD with other splicing components in the activated spliceosome remain to be determined.

CDK11 has additional cellular functions, suggesting that CRK9 shares them, too. In human cells, CDK11 occurs in a mitosis-specific p58 isoform which appears to be involved in mitosis-specific gene expression (65,66). Correspondingly, an initial study on CRK9 provided evidence that the kinase is involved in mitosis and cytokinesis (33). For both kinases though, it has been suggested that their cell cycle-related functions are secondary in nature (15,67). However, tandem affinity purification of CRK9 co-isolated two paralogous, dual-specificity tyrosine phosphorylation-regulated kinases (34) which, due to their predominant locations, were subsequently termed basal body protein 87 (BBP87, Tb927.7.3880) and BBP59 (Tb927.10.350) (68), raising the possibility that CRK9 activity in the nucleus is translocated to the site of cytokinesis initiation by one or both of these kinases. In addition to its splicing function, CDK11 is directly involved in transcription and 3^/^ end processing of human replication-dependent histone genes, which includes CDK11-mediated serine 2 phosphorylation of the heptad repeat sequence YSPTSPS in the C-terminal domain of RPB1, the largest subunit of RNA polymerase II (69). Although conditional *CRK9* silencing led to a loss of RPB1 phosphorylation in trypanosomes (31), it is unlikely that CRK9 shares this function with human CDK11 because histone genes in trypanosomes are part of directional gene arrays that are transcribed polycistronically in replication-independent fashion, and histone pre-mRNAs are polyadenylated (70).

*T. brucei* is transmitted by tsetse flies and causes diseases in humans and livestock animals in sub-Saharan Africa, which are fatal if left untreated. There are no vaccines against the parasite and treatment options are limited. Since CDK11 is an emerging anti-cancer drug target (71–73), CRK9 has the same key function in trypanosome as CDK11 in human gene expression, and the two enzymes have diverged to a degree that renders CRK9 insensitive to the CDK11-selective inhibitor OTS964, it is plausible that a therapeutic drug against *T. brucei* can be developed that selectively inhibits CRK9. This notion is supported by the finding that *CRK9* silencing rescued mice from lethal trypanosome infections, which validated the kinase as a potential chemotherapeutic target in the mammalian host (34). Moreover, it is important to note that in trypanosomatid parasites SL *trans* splicing and polyadenylation are functionally linked (74,75) because polyadenylation has been clinically validated as a chemotherapeutic target in *T. brucei* by the new drug acoziborole which targets the cleavage and polyadenylation specificity factor 3 (76,77). Thus, a CRK9-specific inhibitor may be similarly effective as acoziborole. The quest for such an inhibitor looks promising because the development of a high throughput screening assay for CRK9 seems now within reach, and molecular structure prediction for potential drug candidates has become astonishingly accurate.

## Acknowledgements

We thank Sofia Cassidento, Rakshan Chadha, and Jonathan Hudson for their laboratory assistance and Bruce Mayer for sharing his expertise in kinase biology.

