## supplement for "SF3B1 phosphorylation is an evolutionarily conserved step in spliceosome activation carried out by the divergent, OTS964-insensitive kinase CRK9 in trypanosomes"

**Machida *et al.* – Supplemental material**

**Table S1** OTS964 $EC_{50}$ of procyclic trypanosome viability

**Table S2** OTS964 $IC_{50}$ of CRK9 autophosphorylation *in vitro*

**Table S3** List of DNA oligonucleotides used in RNA analysis

**Figure S1** APmut sequence

**Figure S2** Maps of newly generated plasmids

**Figure S3** Expression in and purification of recombinant GST-SF3B1 proteins from *E. coli*

**Figure S4** SF3B1 amino acid sequence alignment

**Figure S5** OTS964 $EC_{50}$ of human HOP-62 cell viability

**Figure S6** Chai-1-predicted models of the CRK9-OTS964 and $CRK9^{AS}$-OTS964 complexes

***Supplemental References***

**Table S1. Data for OTS964 $EC_{50}$ determination on trypanosome culture growth**

The concentration of OTS964 was serially diluted 1:2 from 250 to 0.12 μM, and the number of viable cells indirectly measured using the luminescent CellTiter-Glo® assay (Promega) according to the manufacturer's specifications.

Lowest and highest average chemiluminescence signals were set to 0 and 100, respectively, and the $EC_{50}$ value calculated by non-linear regression using the GraphPad Prism 10 program

| [OTS964] | chemiluminescence light units | | | | | | | | | | | | | | | | | | | | | | | |
|---|---|---|---|---|---|---|---|---|---|---|---|---|---|---|---|---|---|---|---|---|---|---|---|---|
| | **exp. 1** | **2** | **3** | **4** | **5** | **6** | **7** | **8** | **9** | **10** | **11** | **12** | **13** | **14** | **15** | **16** | **17** | **18** | **19** | **20** | **21** | **22** | **23** | **24** |
| 250 μM | 228 | 198 | 255 | 236 | 238 | 243 | 254 | 253 | 254 | 235 | 264 | 255 | 664 | 333 | 368 | 527 | 659 | 300 | 384 | 522 | 670 | 359 | 403 | 539 |
| 125 μM | 15421 | 12423 | 12231 | 11147 | 15297 | 12548 | 12073 | 10953 | 15198 | 12593 | 12057 | 10974 | 37042 | 31585 | 29857 | 34596 | 37088 | 32564 | 29705 | 35037 | 37395 | 32703 | 30471 | 35541 |
| 62.5 μM | 45823 | 52657 | 50858 | 45152 | 45683 | 52395 | 51092 | 44911 | 45499 | 52007 | 50642 | 45009 | 23630 | 23059 | 40462 | 34521 | 24089 | 23370 | 41231 | 35205 | 24411 | 23625 | 41653 | 35348 |
| 31.25 μM | 24326 | 23458 | 20340 | 20837 | 24606 | 23639 | 20749 | 20832 | 23819 | 23414 | 20687 | 21123 | 28102 | 28313 | 27479 | 26963 | 28497 | 28769 | 28310 | 27531 | 28826 | 28593 | 28460 | 27551 |
| 15.625 μM | 39692 | 38541 | 36482 | 31488 | 39830 | 38928 | 36777 | 31671 | 39593 | 38562 | 36369 | 31750 | 106024 | 66306 | 43237 | 100361 | 107512 | 67508 | 44076 | 102670 | 107919 | 68449 | 44972 | 103280 |
| 7.8125 μM | 277464 | 265226 | 236771 | 274815 | 276036 | 265596 | 237998 | 276636 | 274887 | 266246 | 237533 | 275464 | 477745 | 339866 | 254495 | 457279 | 484973 | 348193 | 261254 | 466770 | 488578 | 351645 | 262672 | 468652 |
| 3.90625 μM | 650450 | 620736 | 578728 | 593066 | 644462 | 618719 | 578587 | 595207 | 636031 | 613400 | 576044 | 592301 | 781271 | 689244 | 577128 | 805472 | 790906 | 701617 | 588907 | 814043 | 791124 | 706459 | 591896 | 815114 |
| 1.953125 μM | 972455 | 945953 | 889346 | 1001511 | 951560 | 932082 | 877859 | 988438 | 925274 | 915752 | 858758 | 962562 | 991578 | 1015815 | 1013981 | 1083869 | 994630 | 1024513 | 1017690 | 1078492 | 981570 | 1013301 | 1014981 | 1065746 |
| 0.976563 μM | 1024115 | 1E+06 | 998814 | 1075963 | 988538 | 977013 | 975351 | 1050005 | 952651 | 943994 | 944110 | 1E+06 | 1033909 | 1078576 | 1117008 | 1061269 | 1029518 | 1078098 | 1114263 | 1056587 | 1E+06 | 1057584 | 1092321 | 1035268 |
| 0.488281 μM | 991156 | 999533 | 912196 | 1038648 | 955362 | 970454 | 888461 | 1007775 | 913823 | 931095 | 855028 | 963984 | 1007466 | 1044844 | 1114728 | 1078970 | 995675 | 1039564 | 1100170 | 1063396 | 968210 | 1019006 | 1075778 | 1035841 |
| 0.244141 μM | 994322 | 955817 | 952478 | 1020684 | 962026 | 927199 | 925245 | 987358 | 921681 | 891781 | 893339 | 942793 | 1063542 | 1056539 | 1096767 | 1059775 | 1048251 | 1044181 | 1078488 | 1040852 | 1E+06 | 1018773 | 1048346 | 1010797 |
| 0.12207 μM | 945434 | 955248 | 917650 | 1019608 | 905820 | 921449 | 887975 | 980938 | 862175 | 881521 | 849059 | 935247 | 1066028 | 1074610 | 1045474 | 1079027 | 1053687 | 1064213 | 1022586 | 1054496 | 1E+06 | 1042159 | 991567 | 1018171 |

| [OTS964] | normalized values | | | | | | | | | | | | | | | | | | | | | | | |
|---|---|---|---|---|---|---|---|---|---|---|---|---|---|---|---|---|---|---|---|---|---|---|---|---|
| 250 μM | -0.013 | -0.016 | -0.010 | -0.012 | -0.012 | -0.011 | -0.010 | -0.010 | -0.010 | -0.012 | -0.009 | -0.010 | 0.030 | -0.003 | 0.001 | 0.016 | 0.029 | -0.006 | 0.002 | 0.016 | 0.030 | 0.000 | 0.004 | 0.017 |
| 125 μM | 1.464 | 1.172 | 1.154 | 1.048 | 1.452 | 1.185 | 1.138 | 1.030 | 1.442 | 1.189 | 1.137 | 1.032 | 3.565 | 3.035 | 2.867 | 3.328 | 3.570 | 3.130 | 2.852 | 3.371 | 3.600 | 3.144 | 2.927 | 3.420 |
| 62.5 μM | 4.419 | 5.083 | 4.908 | 4.354 | 4.405 | 5.058 | 4.931 | 4.330 | 4.387 | 5.020 | 4.887 | 4.340 | 2.262 | 2.206 | 3.898 | 3.320 | 2.306 | 2.237 | 3.973 | 3.387 | 2.338 | 2.261 | 4.014 | 3.401 |
| 31.25 μM | 2.329 | 2.245 | 1.942 | 1.990 | 2.357 | 2.263 | 1.982 | 1.990 | 2.280 | 2.241 | 1.976 | 2.018 | 2.696 | 2.717 | 2.636 | 2.586 | 2.735 | 2.761 | 2.717 | 2.641 | 2.767 | 2.744 | 2.731 | 2.643 |
| 15.625 μM | 3.823 | 3.711 | 3.511 | 3.026 | 3.836 | 3.749 | 3.540 | 3.043 | 3.813 | 3.713 | 3.500 | 3.051 | 10.270 | 6.410 | 4.168 | 9.720 | 10.415 | 6.527 | 4.249 | 9.944 | 10.455 | 6.618 | 4.336 | 10.004 |
| 7.8125 μM | 26.934 | 25.744 | 22.979 | 26.677 | 26.795 | 25.780 | 23.098 | 26.854 | 26.684 | 25.844 | 23.053 | 26.740 | 46.401 | 32.999 | 24.701 | 44.412 | 47.103 | 33.809 | 25.358 | 45.334 | 47.454 | 34.144 | 25.496 | 45.517 |
| 3.90625 μM | 63.188 | 60.299 | 56.216 | 57.610 | 62.606 | 60.103 | 56.203 | 57.818 | 61.786 | 59.586 | 55.955 | 57.536 | 75.903 | 66.958 | 56.061 | 78.255 | 76.840 | 68.161 | 57.206 | 79.088 | 76.861 | 68.632 | 57.496 | 79.193 |
| 1.953125 μM | 94.486 | 91.910 | 86.408 | 97.310 | 92.455 | 90.562 | 85.291 | 96.039 | 89.900 | 88.974 | 83.435 | 93.524 | 96.345 | 98.700 | 98.522 | 105.315 | 96.641 | 99.546 | 98.883 | 104.792 | 95.372 | 98.456 | 98.619 | 103.554 |
| 0.976563 μM | 99.507 | 97.214 | 97.048 | 104.547 | 96.049 | 94.929 | 94.767 | 102.024 | 92.561 | 91.719 | 91.731 | 97.972 | 100.459 | 104.801 | 108.536 | 103.118 | 100.032 | 104.754 | 108.269 | 102.663 | 97.810 | 102.760 | 106.137 | 100.591 |
| 0.488281 μM | 96.304 | 97.118 | 88.629 | 100.920 | 92.824 | 94.291 | 86.322 | 97.919 | 88.787 | 90.466 | 83.072 | 93.662 | 97.889 | 101.522 | 108.315 | 104.839 | 96.743 | 101.009 | 106.900 | 103.325 | 94.073 | 99.011 | 104.529 | 100.647 |
| 0.244141 μM | 96.611 | 92.869 | 92.544 | 99.174 | 93.472 | 90.087 | 89.897 | 95.934 | 89.551 | 86.644 | 86.796 | 91.603 | 103.339 | 102.659 | 106.569 | 102.973 | 101.853 | 101.457 | 104.792 | 101.134 | 99.272 | 98.988 | 101.862 | 98.213 |
| 0.12207 μM | 91.859 | 92.813 | 89.159 | 99.069 | 88.009 | 89.528 | 86.275 | 95.310 | 83.767 | 85.647 | 82.492 | 90.869 | 103.581 | 104.415 | 101.583 | 104.844 | 102.381 | 103.405 | 99.358 | 102.460 | 99.954 | 101.261 | 96.344 | 98.929 |

| | |
|---|---|
| GraphPad Prism 10 | |
| [Inhibitor] vs. normalized response -- Variable slope | |
| Best-fit values | |
| EC50 | 0.000005364 |
| HillSlope | -2.345 |
| logEC50 | -5.271 |
| 95% CI (profile likelihood) | |
| EC50 | 5.194e-006 to 5.540e-006 |
| HillSlope | -2.499 to -2.203 |
| logEC50 | -5.285 to -5.256 |
| Goodness of Fit | |
| Degrees of Freedom | 286 |
| R squared | 0.9815 |
| Sum of Squares | 10242 |
| Sy.x | 5.984 |
| Constraints | |
| EC50 | EC50 > 0 |
| Number of points | |
| # of X values | 288 |
| # Y values analyzed | 288 |

**Table S2. OTS964 $IC_{50}$ determination of CRK9 autophosphorylation**

CRK9 thiophosphate signals, detected with anti-thiophosphoester antibody, was quantified by densitometry using Image Lab Software Version 6.1 (BioRad). For each series, the signal strength of the reaction without inhibitor was set to 100, and $IC_{50}$ calculated by non-linear regression using the GraphPad Prism 10 program.

| [OTS964] | standardized densitometry values | | |
|---|---|---|---|
| | exp.1 | exp. 2 | exp. 3 |
| 0 µM | 100 | 100 | 100 |
| 0.49 µM | 92.934 | 111.048 | 98.825 |
| 0.98 µM | 85.621 | 120.303 | 98.872 |
| 1.95 µM | 90.392 | 122.807 | 95.131 |
| 3.91 µM | 70.614 | 114.682 | 80.472 |
| 7.81 µM | 77.446 | 105.368 | 88.234 |
| 15.63 µM | 72.257 | 89.574 | 92.093 |
| 31.25 µM | 57.657 | 58.136 | 73.476 |
| 62.50 µM | 48.373 | 62.609 | 65.843 |
| 125.00 µM | 37.204 | 45.652 | 54.535 |
| 250.00 µM | 32.038 | 34.698 | 46.334 |
| 500.00 µM | 30.403 | 38.092 | 47.502 |

**Table S3. List of DNA oligonucleotides used in RNA analysis**

| **RNA** | Name | Usage | Sequence (5/ - 3/) |
|---|---|---|---|
| *PAP1* | SLsense PAP1 | 3-primer *trans* splicing assay | **ACAGTTTCTGATCTATATTGGAAGA** |
| | PAP PremRsen1 | 3-primer *trans* splicing assay | **TGATGCACCTTTTTATGTGCTCT** |
| | PAP1 Exon1 Rev | 3-primer *trans* splicing assay | **AATGGAAGGGCATTCGGCCAACA** |
| | Exon 1 Fw | 2-primer *cis* splicing assay | **GTGCAGCGGCACTCCCAAAAC** |
| | Exon 2 Rv | 2-primer *cis* splicing assay | **CGTTAAAACAGATGGACAAATC** |
| α tubulin (*ATUB*) | ATUB Fw q3 | RT-qPCR | **GTGCATTGAACGTGGATCTG** |
| | ATUB Rv q3 | RT-qPCR | **GAGAGTTGCTCGTGGTAGGC** |
| SL RNA | bio-SL_PE | Primer extension | **Biotin-CGACCCCACCTTCCAGATTC** |
| U2 snRNA | bio-U2_PE | Primer extension | **Biotin-ACAGGCAACAGTTTTGATCC** |
| | U2 5' | RT-qPCR | **ATATCTTCTCGGCTATTTAGC** |
| | U2_PE | RT-qPCR | **ACAGGCAACAGTTTTGATCC** |
| U5 snRNA | U5_qPCRF | RT-qPCR | **GCATCGCCGTCTCGAC** |
| | U5_PE | RT-qPCR | **CTATTGAAGCAATATCGG** |
| U6 snRNA | U6FqPCR | RT-qPCR | **GGAGCCCTTCGGGGACA** |
| | U6-3'_rv | RT-qPCR | **GCAAAAGCTATATCTCTCGAAG** |

**A**

```
APmut sequence
   CAAGGAATATCTGCCTCTTTGACATATGGCCGATGAAGACGGTTCTGTTGAACCGACGTTAAAAGCGCATCGCT
   GGGAGATGTCCTCCTCCGCATCTGCAGCTCCCACAACATCGCGGTGGGGCGCGGGGGCACCACGGCAGCGTGGT
   GGCGACGCGCCAAAAGTGGTCGTCAACACGGAGTTGTCGGCGTGGCGTGGGCATGCAGCCCCAGCTCCGAGTGC
   GTACGCGATGGATCCTGCGAAGGCACCAACCATGCGTGCCACTGGTGGTGCGGCGCCGGTGCAAGGAGGTCAAG
   CCCCGATGTTTGGTGGGCAGGCTCCGGTATTTGGAGGTCAGGCACCGGCATTCGGTAGTGGAGCGCCCATGTTC
   ACGGGGGCAGCCCCCGCCGCTGGCATGGCCTTTGGTGCAGCTCCGAATTACCAGTTTGAAGGGACAGCACCTGT
   GCAGTCCCACTTTGCGGGGGGTGAGTCACAATCAGCTGTTACGGCTAACATCGGAGCCCAGGCAAGGAAATTGG
   AAATGCAGTGGCGCGTTAAGAATAAGCGTCTTACAGAGGAGTATCTCGACTCGATCCTTCCACCGGAATTTAAG
   CTTGTGGAACCACCAGCCGATTACAACCCACCGCCCACGGAAGAGCCTAACTTCTACGAGCTTGCCAGTAAGTC
   GTTGGATGTCTTTGTTGTGAACCAAAACAGCGATGCGGCAGCTTCTATGACTTATGATATACCAGAAAGTCTGG
   GAGACGGT
```

**B**

```
          M  A  D  E  D  G  S  V  E  P  T  L  K  A  H  R  W  E  M  S
APmut    ATGGCCGATGAAGACGGTTCTGTTGAACCGACGTTAAAAGCGCATCGCTGGGAGATGTCC   60
         ||||||||||||||||||||||||||||||||||||||||||||||||||||||||||||
WT       ATGGCCGATGAAGACGGTTCTGTTGAACCGACGTTAAAAGCGCATCGCTGGGAGATGTCC   60
          M  A  D  E  D  G  S  V  E  P  T  L  K  A  H  R  W  E  M  S

          S  S  A  S  A  A  P  T  T  S  R  W  G  A  G  A  P  R  Q  R
APmut    TCCTCCGCATCTGCAGCTCCCACAACATCGCGGTGGGGCGCGGGGACGCCACGGCAGCGT  120
         |||||||||||||||||||||||||||||||||||||||||||||.|.||||||||||||
WT       TCCTCCGCATCTGCAGCTCCCACAACATCGCGGTGGGGCGCGGGGGCACCACGGCAGCGT  120
          S  S  A  S  A  A  P  T  T  S  R  W  G  A  G  T  P  R  Q  R

          G  G  D  A  P  K  V  V  V  N  T  E  L  S  A  W  R  G  H  A
APmut    GGTGGCGACACCCCAAAAGTGGTCGTCAACACGGAGTTGTCGGCGTGGCGTGGGCATGCA  180
         |||||||||.|.||||||||||||||||||||||||||||||||||||||||||||||||
WT       GGTGGCGACGCGCCAAAAGTGGTCGTCAACACGGAGTTGTCGGCGTGGCGTGGGCATGCA  180
          G  G  D  T  P  K  V  V  V  N  T  E  L  S  A  W  R  G  H  A

          A  P  A  P  S  A  Y  A  M  D  P  A  K  A  P  T  M  R  A  T
APmut    ACACCAACACCGAGTGCGTACGCGATGGATCCTGCGAAGACCCCAACCATGCGTGCCACT  240
         .|.|||.|.||||||||||||||||||||||||||||||.|.||||||||||||||||||
WT       GCCCCAGCTCCGAGTGCGTACGCGATGGATCCTGCGAAGGCACCAACCATGCGTGCCACT  240
          T  P  T  P  S  A  Y  A  M  D  P  A  K  T  P  T  M  R  A  T

          G  G  A  A  P  V  Q  G  G  Q  A  P  M  F  G  G  Q  A  P  V
APmut    GGTGGTGCGACACCGGTGCAAGGAGGTCAAACTCCGATGTTTGGTGGGCAGACACCGGTA  300
         |||||||||.|.||||||||||||||||||.|.||||||||||||||||||.|.||||||
WT       GGTGGTGCGGCGCCGGTGCAAGGAGGTCAAGCCCCGATGTTTGGTGGGCAGGCTCCGGTA  300
          G  G  A  T  P  V  Q  G  G  Q  T  P  M  F  G  G  Q  T  P  V

          F  G  G  Q  A  P  A  F  G  S  G  A  P  M  F  T  G  A  A  P
APmut    TTTGGAGGTCAGACACCGGCATTCGGTAGTGGAACACCCATGTTCACGGGGGCAACACCC  360
         ||||||||||||.||||||||||||||||||||.|.||||||||||||||||||.|.|||
WT       TTTGGAGGTCAGGCACCGGCATTCGGTAGTGGAGCGCCCATGTTCACGGGGGCAGCCCCC  360
          F  G  G  Q  T  P  A  F  G  S  G  T  P  M  F  T  G  A  T  P

          A  A  G  M  A  F  G  A  A  P  N  Y  Q  F  E  G  T  A  P  V
APmut    GCCGCTGGCATGGCCTTTGGTGCAACTCCGAATTACCAGTTTGAAGGGACAACGCCTGTG  420
         ||||||||||||||||||||||||.||||||||||||||||||||||||||.|.||||||
WT       GCCGCTGGCATGGCCTTTGGTGCAGCTCCGAATTACCAGTTTGAAGGGACAGCACCTGTG  420
          A  A  G  M  A  F  G  A  T  P  N  Y  Q  F  E  G  T  T  P  V
```

**Figure S1**. **APmut sequence**. (**A**) The 748 bp-long APmut sequence was synthesized (IDT) and contained the 25 3′-terminal base pairs of the *T. brucei HSP70* genes 2 and 3 intergenic region and the first 723 bp of the SF3B1 coding region in which 13 threonine codons were mutated to alanine codons. (**B**) Sequence alignment of APmut and wild-type SF3B1 coding sequence, comprising the first 420 bp as well as the corresponding amino acid sequences in blue lettering. TP motifs are marked by green lettering and mutated codon and alanine residues by red lettering.

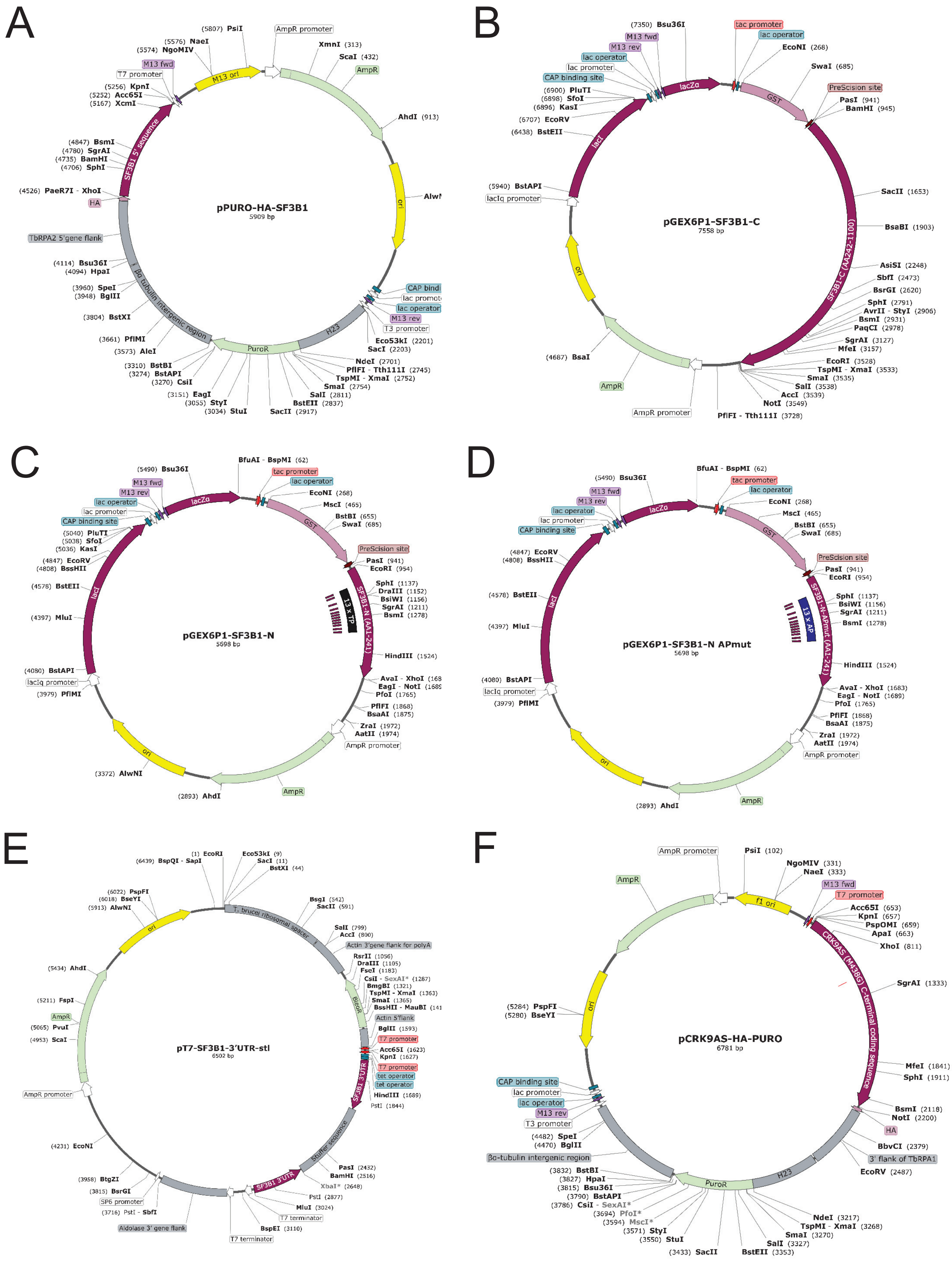
A
pPURO-HA-SF3B1
5909 bp
(5807) PsiI
AmpR promoter
XmnI (313)
ScaI (432)
AmpR
AhdI (913)
ori
(5576) NaeI
(5574) NgoMIV
M13 ori
M13 fwd
T7 promoter
(5256) KpnI
(5252) Acc65I
(5167) XcmI
SF3B1 5' sequence
(4847) BsmI
(4780) SgrAI
(4735) BamHI
(4706) SphI
(4526) PaeR7I - XhoI
HA
TbRPA2 5'gene flank
(4114) Bsu36I
(4094) HpaI
(3960) SpeI
(3948) BglII
(3804) BstXI
βα tubulin intergenic region
(3661) PflMI
(3573) AleI
(3310) BstBI
(3274) BstAPI
(3270) CsiI
(3151) EagI
(3055) StyI
(3034) StuI
PuroR
H23
SacII (2917)
BstEII (2837)
SalI (2811)
SmaI (2754)
TspMI - XmaI (2752)
PflFI - Tth111I (2745)
NdeI (2701)
SacI (2203)
Eco53kI (2201)
T3 promoter
M13 rev
lac operator
B
pGEX6P1-SF3B1-C
7558 bp
(7350) Bsu36I
tac promoter
lac operator
EcoNI (268)
M13 fwd
M13 rev
lac operator
lac promoter
CAP binding site
(6900) PluTI
(6898) SfoI
(6896) KasI
(6707) EcoRV
(6438) BstEII
lacZα
GST
SwaI (685)
PreScission site
PasI (941)
BamHI (945)
lacI
(5940) BstAPI
lacIq promoter
SacII (1653)
BsaBI (1903)
SF3B1-C (AA242-1100)
AsiSI (2248)
SbfI (2473)
BsrGI (2620)
SphI (2791)
AvrII - StyI (2906)
BsmI (2931)
PaqCI (2978)
SgrAI (3127)
MfeI (3157)
EcoRI (3528)
TspMI - XmaI (3533)
SmaI (3535)
SalI (3538)
AccI (3539)
NotI (3549)
PflFI - Tth111I (3728)
AmpR promoter
AmpR
(4687) BsaI
ori
C
pGEX6P1-SF3B1-N
5698 bp
BfuAI - BspMI (62)
(5490) Bsu36I
tac promoter
lac operator
EcoNI (268)
MscI (465)
BstBI (655)
SwaI (685)
M13 fwd
M13 rev
lac operator
lac promoter
CAP binding site
(5040) PluTI
(5038) SfoI
(5036) KasI
(4847) EcoRV
(4808) BssHII
(4578) BstEII
(4397) MluI
(4080) BstAPI
lacIq promoter
(3979) PflMI
lacZα
GST
lacI
PreScision site
PasI (941)
EcoRI (954)
SphI (1137)
DraIII (1152)
BsiWI (1156)
SgrAI (1211)
BsmI (1278)
13 x TP
SF3B1-N (AA1-241)
HindIII (1524)
PfoI (1765)
PflFI (1868)
BsaAI (1875)
ZraI (1972)
AatII (1974)
AmpR promoter
AmpR
(2893) AhdI
ori
(3372) AlwNI
D
pGEX6P1-SF3B1-N APmut
5698 bp
BfuAI - BspMI (62)
(5490) Bsu36I
tac promoter
lac operator
EcoNI (268)
MscI (465)
BstBI (655)
SwaI (685)
M13 fwd
M13 rev
lac operator
lac promoter
CAP binding site
(4847) EcoRV
(4808) BssHII
(4578) BstEII
(4397) MluI
(4080) BstAPI
lacIq promoter
(3979) PflMI
lacZα
GST
lacI
PreScision site
PasI (941)
EcoRI (954)
SphI (1137)
BsiWI (1156)
SgrAI (1211)
BsmI (1278)
13 x AP
SF3B1-N-APmut (AA1-241)
HindIII (1524)
AvaI - XhoI (1683)
EagI - NotI (1689)
PfoI (1765)
PflFI (1868)
BsaAI (1875)
ZraI (1972)
AatII (1974)
AmpR promoter
AmpR
(2893) AhdI
ori
E
pT7-SF3B1-3'UTR-stl
6502 bp
(1) EcoRI
(6439) BspQI - SapI
Eco53kI (9)
SacI (11)
BstXI (44)
(6022) PspFI
(6018) BseYI
(5913) AlwNI
ori
T. brucei ribosomal spacer
BsgI (542)
SacII (591)
SalI (799)
AccI (800)
Actin 3'gene flank for polyA
RsrII (1056)
DraIII (1105)
FseI (1183)
CsiI - SexAI* (1287)
BmgBI (1321)
TspMI - XmaI (1363)
SmaI (1365)
BleoR
Actin 5'flank
BglII (1593)
T7 promoter
Acc65I (1623)
KpnI (1627)
T7 promoter
tet operator
tet operator
HindIII (1689)
PstI (1844)
(5434) AhdI
(5211) FspI
AmpR
(5065) PvuI
(4953) ScaI
AmpR promoter
(4231) EcoNI
(3958) BtgZI
(3815) BsrGI
SP6 promoter
(3716) PstI - SbfI
Aldolase 3' gene flank
T7 terminator
BspEI (3110)
T7 terminator
MluI (3024)
PstI (2877)
XbaI* (2648)
BamHI (2516)
PasI (2432)
stuffer sequence
SF3B1 3'UTR
SF3B1 3'UTR
F
pCRK9AS-HA-PURO
6781 bp
AmpR promoter
PsiI (102)
NgoMIV (331)
NaeI (333)
AmpR
f1 ori
M13 fwd
T7 promoter
Acc65I (653)
KpnI (657)
PspOMI (659)
ApaI (663)
XhoI (811)
SgrAI (1333)
CRK9AS (M438G) C-terminal coding sequence
MfeI (1841)
SphI (1911)
BsmI (2118)
NotI (2200)
HA
BbvCI (2379)
3' flank of TbRPA1
EcoRV (2487)
(5284) PspFI
(5280) BseYI
ori
CAP binding site
lac promoter
lac operator
M13 rev
T3 promoter
(4482) SpeI
(4470) BglII
βα-tubulin intergenic region
(3832) BstBI
(3827) HpaI
(3815) Bsu36I
(3790) BstAPI
(3786) CsiI - SexAI*
(3694) PfoI*
(3594) MscI*
(3571) StyI
(3550) StuI
(3433) SacII
BstEII (3353)
SalI (3327)
SmaI (3270)
TspMI - XmaI (3268)
NdeI (3217)
PuroR
H23

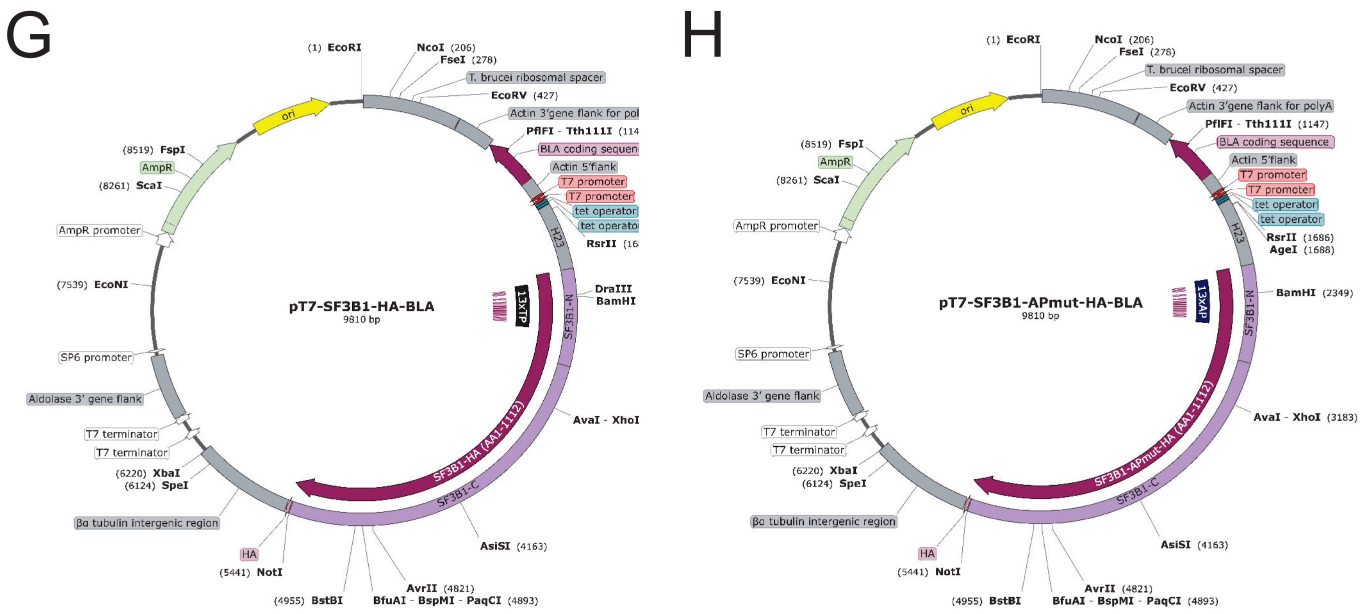

**Figure S2. Maps of newly generated plasmids**. Maps of plasmids PURO-HA-SF3B1 (**A**), GEX6P1-SF3B1-C (**B**), GEX6P1-SF3B1-N (**C**), GEX6P1-SF3B1-N-APmut (**D**), T7-SF3B1-3/UTR-stl (**E**), CRK9$^{AS}$-HA-PURO (**F**), pT7-SF3B1-HA-BLA (**G**) and pT7-SF3B1-APmut-HA-BLA (**H**) were generated with SnapGene software. HA, hemagglutinin tag; H23, intergenic region of *T. brucei HSP70* genes 2 and 3; GST, glutathione-S-transferase tag. BLA, blasticidin S deaminase gene.

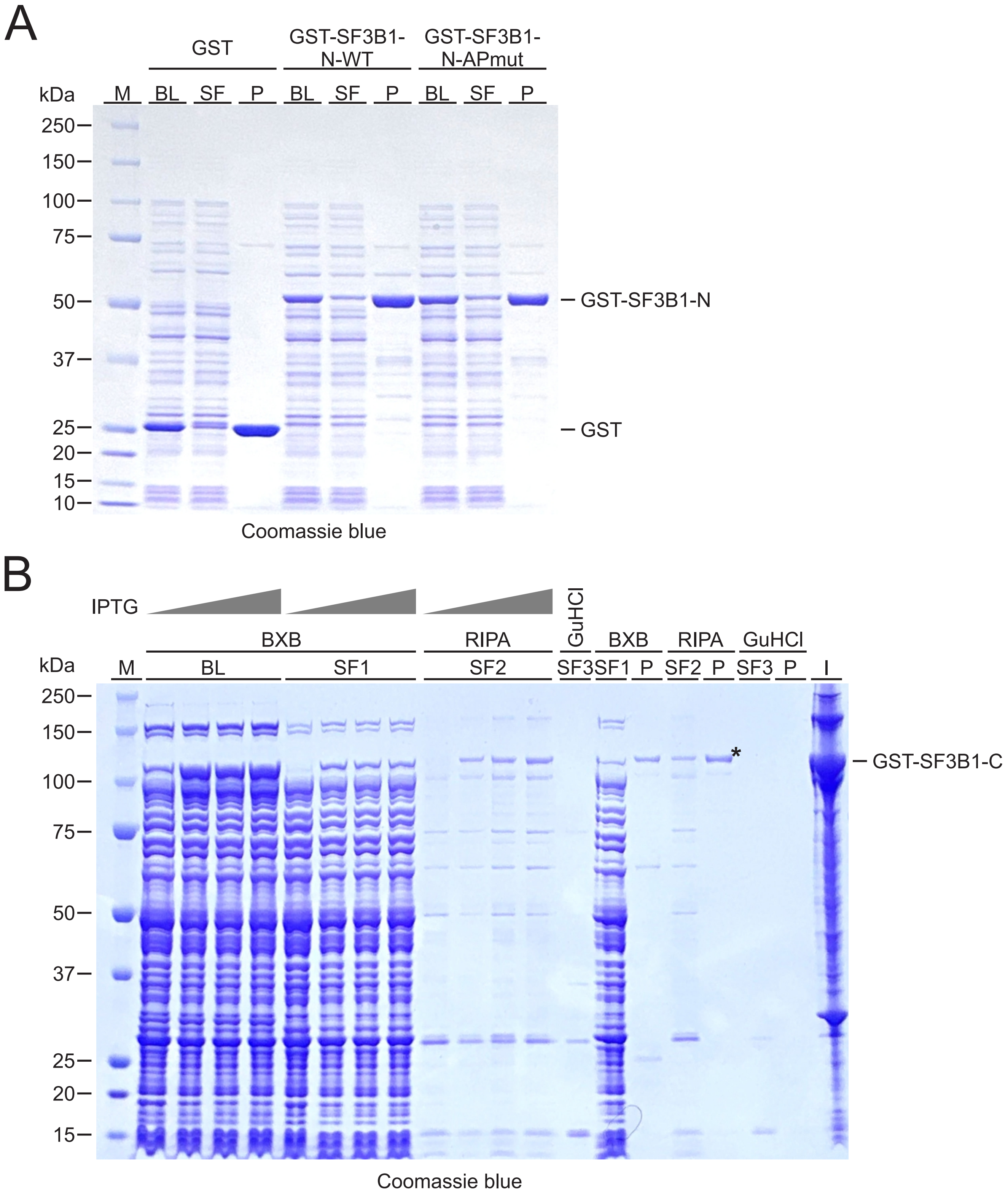

**Figure S3.** **Purification of GST fusion proteins .** (**A**), For crude bacterial lysates (BL), bacteria , expressing GST (26 kDa), GST-SF3B1-N-WT (54 kDa) or GST -SF3B1-N-APmut (54 kDa ) proteins, were dissolved in BXB buffer (PBS containing 1% Triton X-100, 100 mM EDTA, 1mM PMSF, 1% aprotinin). GST fusion proteins were purified from the soluble fraction (SF) with glutathione agarose beads and directly released from pellets (P) into 1x SDS protein loading buffer. ( **B**), Since GST-SF3B1-C (124 kDa) was predominantly insoluble in bacteria cultured at 37°C, the protein was expressed in *E. coli* DH5α at 18°C for 24 hours in the presence of 0.05 - 0.2 mM IPTG. Subsequently, the bacterial pellet was sequentially treated with BXB, RIPA buffer (150 mM NaCl , Tris-HCl pH 8.0, 0.1% SDS, 1% sodium deoxycholate, 5 mM EDTA) and denaturing buffer (2 M guanidine hydrochloride [GuHCl], 200 mM NaCl, 10 mM sodium phosphate). Soluble fractions are indicated as SF1-3. For *in vitro* kinase assays, GST-SF3B1-C purified with RIPA buffer was used (lane RIPA/P, see asterisk). I, insoluble fraction after BXB treatment.

```
HsSF3B1    MAKIAKTHEDIEAQIREIQGKKAALDEAQGVGLDSTGYYDQEIYGGSDSRFAGYVTSIAA   60
HsSF3B1    TELEDDDDDYSSSTSLLGQKKPGYHAPVALLNDIPQSTEQYDPFAEHRPPKIADREDEYK  120
HsSF3B1    KHRRTMIISPERLDPFADGGKTPDPKMNARTYMDVMREQHLTKEEREIRQQLAEKAKAGE  180

HsSF3B1    LKVVNGAAASQPPSKRKRRWDQTADQTPGATPKKLSSWDQAETPGHTPSLRWDETPGRAK  240
TbSF3B1    ---MADEDGSVEPTLKAHRWEMSS------SA-----------SAAPTTSRWGAGTPRQR   40
LmSF3B1    ---MADAEGGEPQALKKHRWEDPTGFGGGDSG-----------GAVPTSTRWSSATPRQY   46
              : .  ..   : : :**:  :      :             .   : **.    *

HsSF3B1    GSETPGATPG----SKIWDPTPSHTPAGA-----ATPGRGDTPGHATPGHGGATSSARKN  291
TbSF3B1    GGDTPKV--VVNTELSAWRGHATPTPSAYAMDPAKTPTMRAT---GG-----ATPV----   86
LmSF3B1    GADTPRRTPVLSSETSSWRGMATPNATPYMTDFIPTPTFGAS---GSSGMGGSTPSTFGT  103
           *.:**          . *    : . :        **    :   .      :*

HsSF3B1    R--WDETPKTERDTPGHGSGWAETPRTDRGG-DSIGETPTPGASKRKSRWDETPASQMGG  348
TbSF3B1    --QGGQTPMFGGQTPVF-GG--QTPAFGSGTPMFTGATPAAG-----MAFGATPNYQFEG  136
LmSF3B1    PRMGSSGVAAGGSTPLFASG--STPRMGGVTPMFTGATPAAG-----ATFGATPNYQYEG  156
               ..      .** . .*  .**  .       * **: *       :. **  *  *

HsSF3B1    STPVLTPGKTPIGTPAMNMATPTPGHIMSMTPEQLQAWRWEREIDERNRPLSDEELDAMF  408
TbSF3B1    TTPVQSHFAGGE-S-----------QSAVTANIGAQARKLEMQWRVKNKRLTEEYLDSIL  184
LmSF3B1    GTPLQSHFASSVET-----------QSITTAAIEAKAKALEKEWNRKNKRLTSEYLDSIL  205
            **: :       :           :    :    :*   * :   :*: *:.* **:::

HsSF3B1    PEGYKV-LPPPAGYVPIRTPARKLTATPTPLGGMTGFHMQTEDRT-------------MK  454
TbSF3B1    PPEFKL-VEPPADYNPPPTEEPNFYEL--ASKSLDVFVVNQNSDA---------AASMTY  232
LmSF3B1    PREFFTVAAVPADYNPLPSEEPNFYEI--AVRSMDVFSMQAGAAAAVTTGLDANGQPIRY  263
           *  :      **.* *  :   ::        .:  * ::    :

HsSF3B1    SVNDQPSGNLPFLKPDDIQYFDKLLVDVDESTLSPEEQKERKIMKLLLKIKNGTPPMRK  514
TbSF3B1    DIPESLGDGMPQMQTQDAPVFEVLLKYHNVNPIPDEVLPSYMLMKNLFKIKNGDTNQRRI  292
LmSF3B1    DIPEDMGDGMPAMKVEDAAVFVELLKYHKAEKIPDEHLPSYLLMKNLFKIKNGDTMQRR  323
           .: :. ...:* :: :*   *  **   . . :  *   .  :** *:*****    *:

                         xxxxxxxx HEAT 1 xxxxxxxxxxxxxxxxxxxxxxxxxxxxxx
HsSF3B1    ALRQITDKAREFGAGPLFNQILPLLMSPTLEDQERHLLVKVIDRILYKLDDLVRPYVHKI  574
TbSF3B1    AMRYLLDKARIFGSGPLFQFTFHVWRSGILDVIEQHYYVDLVKGAISRLQKDVRGSSKEI  352
LmSF3B1    GTRYLLDKATIFGSHFIFQRLSYIWSSDILNIEEKHYFIDFIKSLLQQMGKGARQFTKEV  383
           . * : ***  **:  :*:    :  *  *:  *:*  :..:.  : :: . .*   :::

           xx HEAT 2 xxxxxxxxxxxxxxxxxxxxxxxxxxx HEAT 3 xxxxxxxxxxxxxxx
HsSF3B1    LVVIEPLLIDEDYYARVEGREIISNLAKAAGLATMISTMRPDIDNMDEYVRNTTARAFAV  634
TbSF3B1    VHMMEVLLSAQERVLRDDGKEVLILLTRVVGYEAVFEAIKEDFAHAESGVRRHTAKVVAI  412
LmSF3B1    IHLVEPLLTAHERILRDDGKQVLTLLTRVVGFQAVFAVIREDFGHAESIVRRHTARVMAV  443
           : ::* **  .:   * :*::::  *::..*  ::: .:: *: : :. **. **:..*:

           Xxxxxxx xxxxxxxx HEAT 4 xxxxxxxxxxxxxxxxxxx  xxxxx HEAT 5 xx
HsSF3B1    VASALGIPSLLPFLKAVCKSKKSWQARHTGIKIVQQIAILMGCAILPHLRSLVEIIEHGL  694
TbSF3B1    IAFAVGPEIAIKIIHGMSLSP-VALARQTSARAITEMATILMHAITGELVELVPIFEKLL  471
LmSF3B1    VGYAAGTEEVTAALRDMSYAP-SALARQTVVRSMAELAKMVGHALIAALPEMVAMLERLL  502
           :. * *       :: :. :     **:*  : : ::* ::  *:   * .:* ::*: *
           xxxxxxxxxxxxxxxxxxxxxxxx
HsSF3B1    VDEQQKVRTISALAIAALAEAATPYGIESFDSVLKPLWKGIRQHRGKGLAAFLKAIGYLI  754
TbSF3B1    RDE-PRVKREAASALAHVAEATYPYGIEELDSLVYIVREECKRGIGTTTGLFVRAFGALI  530
LmSF3B1    RDE-KRVQRDAALAVAAIAEATAPDGIEELAPLVDVICEECMKGIGSMASPFIQAFGALV  561
            **  :*:  :* *:* :***: * ***.:  ::  : :   :  *.  . *::*:* *:

                   xxxxxxxx HEAT6 xxxxxxxxxxxxxxxxxxxxxxxx
HsSF3B1    PLMDAEYANYYTREVMLILIREFQSPDEEMKKIVLKVVKQCCGTDGVEANYIKTEILPPF  814
TbSF3B1    PLMAPYDAQKYTSDMLPTLVNQFNTPDDEHRRVLLQVVRQCVSADGVTVDFIRNVILRPF  590
LmSF3B1    PLMSPYDAQARTAAMMPNLVNQFSTPEDEFRRVLISVVRKCVLAEGVTPQFIRQTILEPF  621
           ***    *:  *  ::  *:.:*.:*::* :::::.**::*  ::**  ::*:  ** **

                                        xxxxxxxxxxxx HEAT 7 xxxxxxxxxxx
HsSF3B1    FKHFWQHR-MALDRRNYRQLVDTTVELANKVGAAEIISRIVDDLKDEAEQYRKMVMETIE  873
TbSF3B1    FEGFWTIRRVAADRKTAGILIDTTVGIARRIGSTEILQYLVQDMKDENEHFQRMVVETVR  650
LmSF3B1    FEGFWRVRRLAAERQTSGSLVATTVEIAKKLGSVDVLVKLSPDMKDESEEYQRMVLTAIK  681
           *: **  * :* :*:.   *: *** :*.::*:.:::  :  *:*** *.:::**: ::.
```

```
          xxxxxxxx
HsSF3B1   KIMGNLGAADIDHKLEEQLIDGILYAFQ-EQTTEDSVMLNGFGTVVNALGKRVKPYLPQI 932
TbSF3B1   RVIANVGAVGVPDTLVALLMDGAIAAVKQDETGLNRVVMEGLATICNALGTRLKRHLRQI 710
LmSF3B1   KVVDATGMDAAPDTLVTFVLDGAIAAVRQDELGTSKLVMNVLATICNALAARLRPYLKQV 741
          :::   *     ..*   ::** : *.: ::   . :::: :.*: ***. *:: :* *:

HsSF3B1   CGTVLWRLNNKSAKVRQQAADLISRTAVVMKTCQEEKLMGHLGVVLYEYLGEEYPEVLGS 992
TbSF3B1   FDLIKSRRDM-PGMIRMQAAELAARIATTVKDAGGALFLQDLGRSLFDRLEDDEAAVMSA 769
LmSF3B1   FDLIKRRRENREASMRAQTADLVSRIARTVMLADGAVFLQDLGLSLYERLEDPDARALSA 801
           . :  * :   . :* *:*:* :* * .:  .    :: .**  *:: * :    .:.:

                           xxxxxxxx HEAT 8 xxxxxxxxxxxxxxxxxxxxxxx   x
HsSF3B1   ILGALKAIVNVIGMHKMTPPIKDLLPRLTPILKNRHEKVQENCIDLVGRIADRGAEYVSA 1052
TbSF3B1   NLKATRVILVELGAARYQPPVRELLKKLMYIIPNRNSNVQLNTILLVEEIATNCDEDVEA 829
LmSF3B1   NLRAVRSILGELGSRRFKPSVRELLKRLTFVIKSRNSHVQNSAIALIEDIATNYDTDVDA 861
           * * : *:  :*  :  * :::** :*  :: .*:.:** . * *:  ** .    *.*

          xxxxxx-x HEAT 9 xxxxxxxxxxxxxxxxxxxxxxx
HsSF3B1   REWMRI-CFELLELLKAHKKAIRRATVNTFGYIAKAIGPHDVLATLLNNLKVQERQNRVC 1111
TbSF3B1   IHLQELATKGLFELLDAHRRETRRACTRTFGVIARKIRPFAIILELVDNFKQDKRKIRIC 889
LmSF3B1   IHLHQLATRGLFELLDSPQRATRHACARTFGVIAKKIRPFAIILELVDNFRQDKRQIRIC 921
           .  .:    *:***.: ::  *:* ..*** **: * *. ::  *::*:: ::*: *:*

                    xxxxxxx HEAT 10 xxx-----xxxxxxxxxxxxxxxxxxxx  xxxx
HsSF3B1   TTVAIAIVAETCSPFTVLPALMNEYRVPE-----LNVQNGVLKSLSFLFEYIGEMGKDYI 1166
TbSF3B1   TAVALGAIARECGAFTVIPYLLNESKICEGEQVATIVQHSILKAVRYIFEAIGAAGKDFV 949
LmSF3B1   TAVALSAIAKECGPFTIIPYLLNEYKISEGKQVAVIVQHSVLKAIRYIFEAIGSIGKEYV 981
          *:**:. :*. *. **::* *:** :: *       **:.:**:: ::** **  **:::

          xxxx HEAT 11 xxxxxxxxxxxxxxxxxxxxxx
HsSF3B1   YAVTPLLEDALMDRDLVHRQTASAVVQHMSLGVYG-FGCEDSLNHLLNYVWPNVFETS-- 1223
TbSF3B1   YPLVPLLVRALTEMEIQHRRMAVEACRSIVLAVAGNDGFEDLVIHFLNFIHPNIVELLSR 1009
LmSF3B1   YPMIPLLERALTETNIQMRRMAVEACRAILLSVAGNDGFEDIALHLLNFVHPNIVELLAK 1041
          * : ***  ** : ::  *: *  . : : *.* *  * **   *:**:: **:.*

                                           =SF3B3 & SF3B5 interaction=
HsSF3B1   ---------PHVIQAVMGALEGLRVAIGPCRMLQYCLQGLFHPARKVRDVYWKIYNSIYI 1274
TbSF3B1   NETKISEERLKMVTAVVGYYEAARLVIGSGKLFQYLLQGLFHPAKKVRDIYRRTYNMVYM 1069
LmSF3B1   NEVKIGEERLKMVTAVVSYYEAARLVIEPGKLLQYLLQGLFHPARKVRDIYRRTYNLIYV 1101
                    ::: **:.  *. *:.*   :::** ********:****:* : ** :*:

          ==============================
HsSF3B1   GSQDALIAHYPRIYNDDKNTYIRYELDYIL    1304
TbSF3B1   ASPEALVPYYPRLGDDNEHTYVRHELEVLL    1099
LmSF3B1   GSPERLVPYYPRIENDASHTYVRHELEVLL    1131
          .* : *: :***: :* .:**:*:**: :*
```

**Figure S4. SF3B1 amino acid sequence alignment.** SF3B1 sequences of *Homo sapiens* (*Hs*; accession number NP_036565/UniProt O75533), *Trypanosoma brucei* (*Tb;* Tb927.11.11850/Q382Z6) and *Leishmania major* (*Lm*; LmjF.28.2570/Q4Q804) were aligned using CLUSTAL Omega (v1.2.4) with default parameters (1,2). Blue and black lettering indicate the N- and C-terminal domains of SF3B1, respectively. TP and SP motifs, potential CDK phosphorylation motifs, are highlighted in yellow. The HEAT domains, as annotated for human SF3B1 at UniProt are outlined above the sequences. Asterisks, colons and dots indicate identical, conserved and semi-conserved positions among the three sequences, respectively.

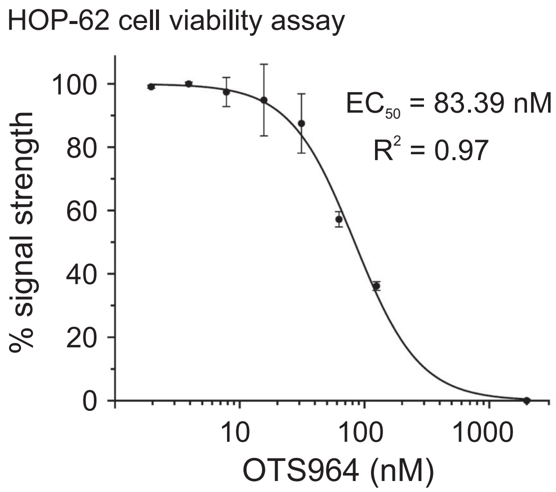

**Figure S5.** **$EC_{50}$ of OTS964 on human HOP-62 cell viability.** The HOP-62 human non-small cell lung cancer cell line was cultured in RPMI 6226 medium supplemented with 10% fetal bovine serum. 1x $10^5$ cells were trypsinized and plated on 96 well dishes in the presence or absence of OTS964 concentrations ranging from 1 nM to 2 μM. Cell viability was measured after 48 h using the CellTiter-Glo Luminescent Cell Viability Assay (Promega) with luminescence detection on a TECAN SPARK microplate reader. $EC_{50}$ was determined by non-linear curve fitting using the GraphPad Prism program (v10. 5.0) and calculated to be 83.39 nM.

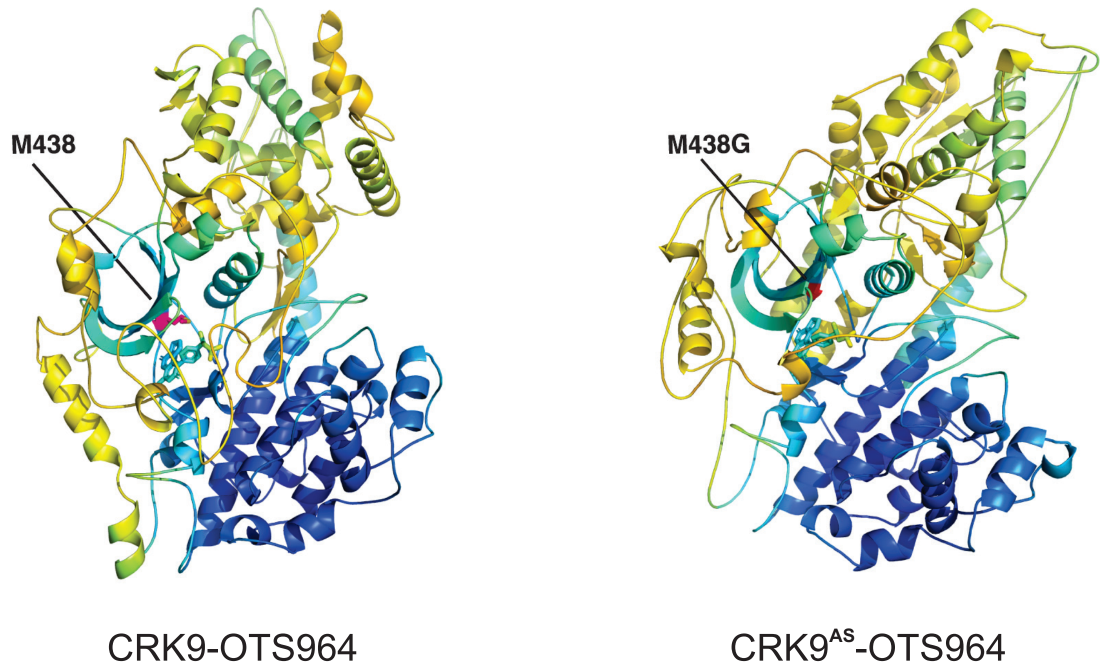

**Figure S6.** **Chai-1-predicted models of the CRK9-OTS964 and CRK9AS-OTS964 complexes**. OTS964 and M438 (gatekeeper residue) are shown as licorice sticks. The predicted local distance difference test (pLDDT) scores are mapped onto the models, with blue indicating regions of high confidence and yellow to red indicating regions of lower confidence. The predicted template modeling (pTM) and interface predicted template modeling (ipTM) scores are 0.63 and 0.59 for CRK9, and 0.59 and 0.59 for CRK9AS, respectively.

### *Supplemental References*

1. Madeira, F., Madhusoodanan, N., Lee, J., Eusebi, A., Niewielska, A., Tivey, A.R.N., Lopez, R. and Butcher, S. (2024) The EMBL-EBI Job Dispatcher sequence analysis tools framework in 2024. *Nucleic Acids Res*, **52**, W521-W525.

2. Sievers, F., Wilm, A., Dineen, D., Gibson, T.J., Karplus, K., Li, W., Lopez, R., McWilliam, H., Remmert, M., Soding, J. *et al.* (2011) Fast, scalable generation of high-quality protein multiple sequence alignments using Clustal Omega. *Mol. Syst. Biol.*, **7**, 539.
